# Adaptive diversification through structural variation in barley

**DOI:** 10.1101/2024.02.14.580266

**Authors:** Murukarthick Jayakodi, Qiongxian Lu, Hélène Pidon, M. Timothy Rabanus-Wallace, Micha Bayer, Thomas Lux, Yu Guo, Benjamin Jaegle, Ana Badea, Wubishet Bekele, Gurcharn S. Brar, Katarzyna Braune, Boyke Bunk, Kenneth J. Chalmers, Brett Chapman, Morten Egevang Jørgensen, Jia-Wu Feng, Manuel Feser, Anne Fiebig, Heidrun Gundlach, Wenbin Guo, Georg Haberer, Mats Hansson, Axel Himmelbach, Iris Hoffie, Robert E. Hoffie, Haifei Hu, Sachiko Isobe, Patrick König, Sandip M. Kale, Nadia Kamal, Gabriel Keeble-Gagnère, Beat Keller, Manuela Knauft, Ravi Koppolu, Simon G. Krattinger, Jochen Kumlehn, Peter Langridge, Chengdao Li, Marina P. Marone, Andreas Maurer, Klaus F.X. Mayer, Michael Melzer, Gary J. Muehlbauer, Emiko Murozuka, Sudharsan Padmarasu, Dragan Perovic, Klaus Pillen, Pierre A. Pin, Curtis J. Pozniak, Luke Ramsay, Pai Rosager Pedas, Twan Rutten, Shun Sakuma, Kazuhiro Sato, Danuta Schüler, Thomas Schmutzer, Uwe Scholz, Miriam Schreiber, Kenta Shirasawa, Craig Simpson, Birgitte Skadhauge, Manuel Spannagl, Brian J. Steffenson, Hanne C. Thomsen, Josquin F. Tibbits, Martin Toft Simmelsgaard Nielsen, Corinna Trautewig, Dominique Vequaud, Cynthia Voss, Penghao Wang, Robbie Waugh, Sharon Westcott, Magnus Wohlfahrt Rasmussen, Runxuan Zhang, Xiao-Qi Zhang, Thomas Wicker, Christoph Dockter, Martin Mascher, Nils Stein

## Abstract

Pangenomes are collections of annotated genome sequences of multiple individuals of a species. The structural variants uncovered by these datasets are a major asset to genetic analysis in crop plants. Here, we report a pangenome of barley comprising long-read sequence assemblies of 76 wild and domesticated genomes and short-read sequence data of 1,315 genotypes. An expanded catalogue of sequence variation in the crop includes structurally complex loci that have become hot spots of gene copy number variation in evolutionarily recent times. To demonstrate the utility of the pangenome, we focus on four loci involved in disease resistance, plant architecture, nutrient release, and trichome development. Novel allelic variation at a powdery mildew resistance locus and population-specific copy number gains in a regulator of vegetative branching were found. Expansion of a family of starch-cleaving enzymes in elite malting barleys was linked to shifts in enzymatic activity in micro-malting trials. Deletion of an enhancer motif is likely to change the developmental trajectory of the hairy appendages on barley grains. Our findings indicate that rapid evolution at structurally complex loci may have helped crop plants adapt to new selective regimes in agricultural ecosystems.

---

Reliable crop yields fueled the rise of human civilizations. As people embraced a new way of life, cultivated plants, too, had to adapt to the needs of their domesticators. There are different adaptive requirements in a wild compared to an arable habitat. Crop plants and their wild progenitors differ in how many vegetative branches they initiate or how many seeds or fruits they produce and when. For example, barley (*Hordeum vulgare*) in six-rowed forms of the crops, thrice as many grains set as in the ancestral two-rowed forms. This change was brought about by knock-out mutations^1^ of a recently evolved regulator^2^ of inflorescence development. Consequently, six-rowed barleys came to predominate in most barley-growing regions^3^. Taking a broader view of the environment as a set of exogeneous factors that drive natural selection, barley provides another fascinating, and economically important example. The process of malting involves the sprouting of moist barley grains, driving the release of enzymes that break down starch into fermentable sugars. In the wild, various environmental cues can trigger germination to improve the odds of the emerging seedling encountering favorable weather conditions for subsequent growth^4^. In the malt house, by contrast, germination of modern varieties has to be fast and uniform to satisfy the desired specifications of the industry. In addition to these examples, traits such as disease resistance, plant architecture and nutrient use have been both a focus for plant breeders and studied intensively barley geneticists^5^. While barley genetic analyses flourished during a “classical” period^6^ in the first half of the 20th century, it started to lag behind small-genome models due to difficulties in adapting molecular biology techniques to a large genome rich in repeats^7^. However, interest in barley as diploid model for temperate cereals has surged again as DNA sequencing became more powerful. High-quality sequences of several barley genomes have been recently assembled^8^. New sequencing technologies have shifted the focus of barley genomics: from the modest ambition of a physical map of all genes to a “pangenome”, i.e. near-complete sequence assemblies^9^ of many genomes. Here, we report a pangenome comprising 76 chromosome-scale sequences assembled from long-reads as well as short-read sequences of 1,315 barley genomes. These data in conjunction with genetic and genomic analyses provide insights into the effects of structural variation at loci related to crop evolution and adaptation.

## An expanded annotated pangenome of barley

As in previous diversity studies^8,10^, we aimed for a judicious mix of representativeness, diversity and integration with community resources (**Fig. 1a**, **Extended Data** Fig. 1a-c, **Supplementary Table 1**). We selected (i) diverse domesticated germplasm with a focus on genebank accessions from barley’s center of diversity in the Middle East; (ii) 23 accessions of barley’s conspecific wild progenitor *H. vulgare* subsp. *spontaneum* from across that taxon’s geographic range (**Extended Data** Fig. 1d); and (iii) cultivars of agronomic or scientific relevance. Examples of the last category are Bonus, Foma and Bowman, three parents of classical mutants^11^. Genome sequences of each accession were assembled to contig-level from PacBio HiFi accurate long reads^12^ and scaffolded with conformation capture sequencing (Hi-C) data^13^ to chromosome-scale pseudomolecules (**Extended Data** Fig. 2a, **Supplementary Table 1)**. Gene models were annotated with the help of transcriptional evidence and homology. Illumina RNA sequencing and PacBio isoform sequencing of five different tissues (**Supplementary Table 2**) were generated for 20 accessions. Gene models predicted in these genomes were projected onto the remaining 56 sequence assemblies (**Supplementary Table 3**). Out of 4,896 single-copy genes conserved across the Poales, on average fewer than 92 (1.9%) were absent in the pangenome annotations (**Supplementary Table 3**). Our assemblies also met the other quality metrics proposed by the EarthBiogenome project^14^ (**Supplementary Table 1**).

**Figure 1:**
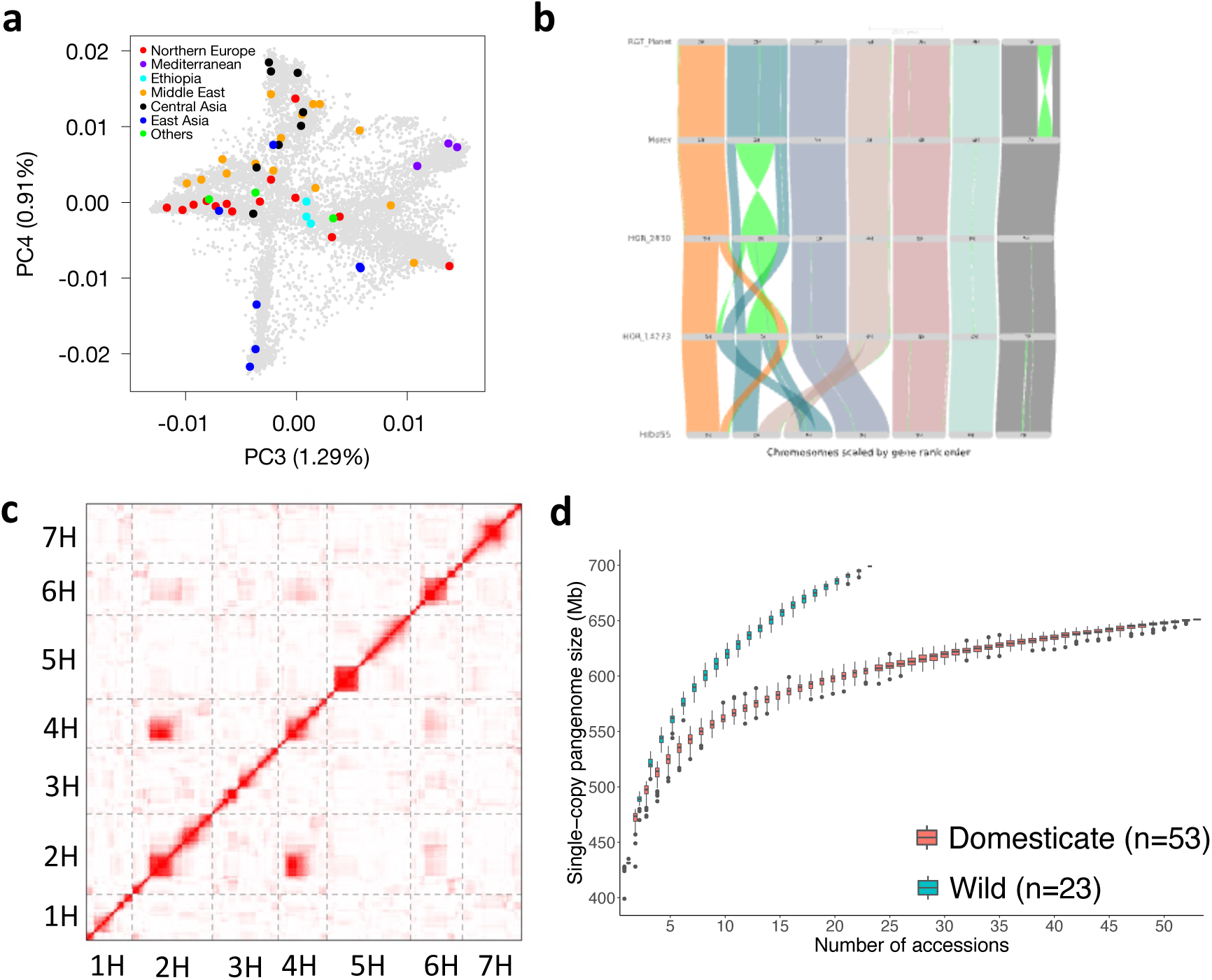
A species-wide pangenome of *Hordeum vulgare.* **(a)** Principal component analysis showing domesticated accessions (n=53) in the pangenome panel in the global diversity space. Regions of origins are color coded. The proportion of variance explained by each PC in panels is given in the axis labels. Other PCs are shown in **Extended Data** Fig. 1a. **(b)** Example of large structural variants including interchromosomal translocations and inversions between pangenome accessions. **(c)** Interchromosomal linkage disequilibrium (LD) in segregating offspring derived from a cross between HID055 and Barke. LD is indicated by the intensity of red color. **(d)** Size of the single-copy pangenome in wild and domesticated barleys as a function of sample size.

## An atlas of structural variation

Gene content variation was abundant in the barley pangenome. The gene models in the 76 genomes were clustered into 95,735 orthologous groups (**Extended Data** Fig. 3), of which only 16,672 (17.4%) were present in all 76 genotypes. Of these groups, 14,736 had a single representative in each of the genomes. At the level of individual gene models, a third were considered conserved because they belong to an orthologous group with representatives from each accession (**Extended Data** Fig. 3b). As expected for conspecific populations connected by gene flow, wild and domesticated barleys were not strongly differentiated in their gene content: of 78,565 orthologous groups subject to presence/absence variation, only 863 and 397 were private to wild and domesticated barleys, respectively. The functional annotations of clusters restricted to specific gene pools (wild forms, landraces, cultivars and combinations of these groups) pointed to an involvement in biotic and abiotic stress responses (**Supplementary Table 4**).

To expand the catalogue of presence/absence variants (PAV), insertion and deletions (indels) and polymorphic inversions, we aligned the genome sequences and detected structural variants (**Fig. 1b**, **Extended Data** Fig. 2b-d, **Extended Data** Fig. 3c). Noteworthy were two reciprocal interchromosomal translocations, the first in HOR 14273, an Iranian landrace, and the second in HID055, a wild barley from Turkey (**Fig. 1b**). The latter event joins the short arm of chromosome 2H with the long arm of chromosome 4H (and vice versa) and manifests itself in a biparental population between HID055 and Barke^15^ in interchromosomal linkage (**Fig. 1c**) and incomplete seed set in the offspring. This illustrates that inadvertent selection of germplasm with structural variants can create obstacles for the use of plant genetic resources. The presence of both wild and domesticated barleys in our panel made it possible to compare the levels of structural diversity in the two taxa. Graph structures tabulating the presence and absence of single-copy loci in individual genomes^8^ grew faster in wild than in cultivated forms (**Fig. 1d**): a larger amount of single-copy sequence was present in 23 wild barley genomes than in 53 genomes of the domesticate. This pattern was also seen in a whole-genome graph constructed with minigraph^16^ (**Extended Data** Fig. 4e). The pangenome graph improves the accuracy of read alignment and variant calling: more reads were aligned as proper pairs, and with fewer mismatches, to the graph than to a single reference genome (**Extended Data** Fig. 4b). The genome-wide distribution of structural variants encapsulated in the graph matched that inferred from pairwise alignments (**Extended Data** Fig. 4c-d). However, owing to high computational requirements^17^, pangenome graph construction with packages supporting small variants (< 50 bp) is still computationally prohibitive in barley.

Despite domestication bottlenecks, genetic diversity is high in cultivated barley^5^. To quantify the completeness of the haplotype inventory of our pangenome, we compared our assemblies against short-read data of a global diversity panel (**Supplementary Table 5**). A core set of 1,000 genotypes selected from a collection of 22,626 barleys^3^ was sequenced to three-fold haploid genome coverage. Nested therein, 200 genomes^8^ were sequenced to 10-fold depth and the gene space of 46 accessions was represented in the contigs assembled from 50-fold short-read data (**Extended Data** Fig. 5a**, Supplementary Table 6)**. A total of 315 elite cultivars of European ancestry were sequenced to 3-fold coverage (**Extended Data** Fig. 5a**, Supplementary Table 5**). More than 164.5 million single-nucleotide polymorphisms (SNPs) and indels were detected across all panels (**Extended Data** Fig. 5b). Overlaying these with the pangenome showed that the 76 chromosome-scale assemblies captured almost all pericentric haplotypes of cultivated barley (**Extended Data** Fig. 2d-f). Coverage decreased to as low as 50% in distal regions, where haplotypes of plant genetic resources lacked a close relative in the pangenome more often than those of elite cultivars (**Extended Data** Fig. 2e-f). This suggests that, thanks to broad taxon sampling, short-read sequencing will remain indispensable for the time being, but in the future population-scale long-read sequencing^18^ will be a desirable in agricultural genetics as it is in medical genetics.

## An inventory of complex loci

Long-read sequencing has the power to resolve structurally complex genomic regions, where repeated cycles of tandem duplication, mutation of duplicated genes and elimination by deletion or recombination have created a panoply of diverged copies of one or multiple genes in varied arrangements (**Extended Data** Fig. 6a). Many complex loci are intimately linked to the evolution of resistance genes^19^. An illustrative example is barley’s *Mildew resistance locus a (Mla)*^20,21^, which contains three families of resistance gene homologs, each with multiple members at the locus. A 40 kb region containing members of two families is repeated four times head-to-tail in RGT Planet, but is not present in even a single complete copy in 62 accessions of our pangenome (**Extended Data** Fig. 6b-c). *Mla* genes *sensu strictu*, i.e. those that have been experimentally proven to provide functional powdery mildew resistance, are among members of a subfamily that resides outside of this duplication but close to its distal border (**Fig. 2a-b**, **Extended Data** Fig. 6b-c). Twenty-nine *Mla* alleles in the narrow sense have been defined to date^22^. Gene models identical to seven were identified in our pangenome (**Fig. 2a**). However, the sequence variation went beyond this observation: 149 unique gene models were different from, but highly similar to known *Mla* alleles, with nucleotide sequences at least 98% identical. Some of these genes were present in multiple copies. HOR 8117, a landrace from Nepal, contained 11 different close homologs of *Mla,* two of which were present in five copies each (**Supplementary** Fig. 1). Genome sequences alone cannot inform us how this sequence diversity relates to resistance to powdery mildew or other diseases^23^. Until the advent of long-read sequencing, it was virtually impossible to resolve the structure of the *Mla* locus in multiple genomes at once, but now it is a corollary of pangenomics.

**Figure 2:**
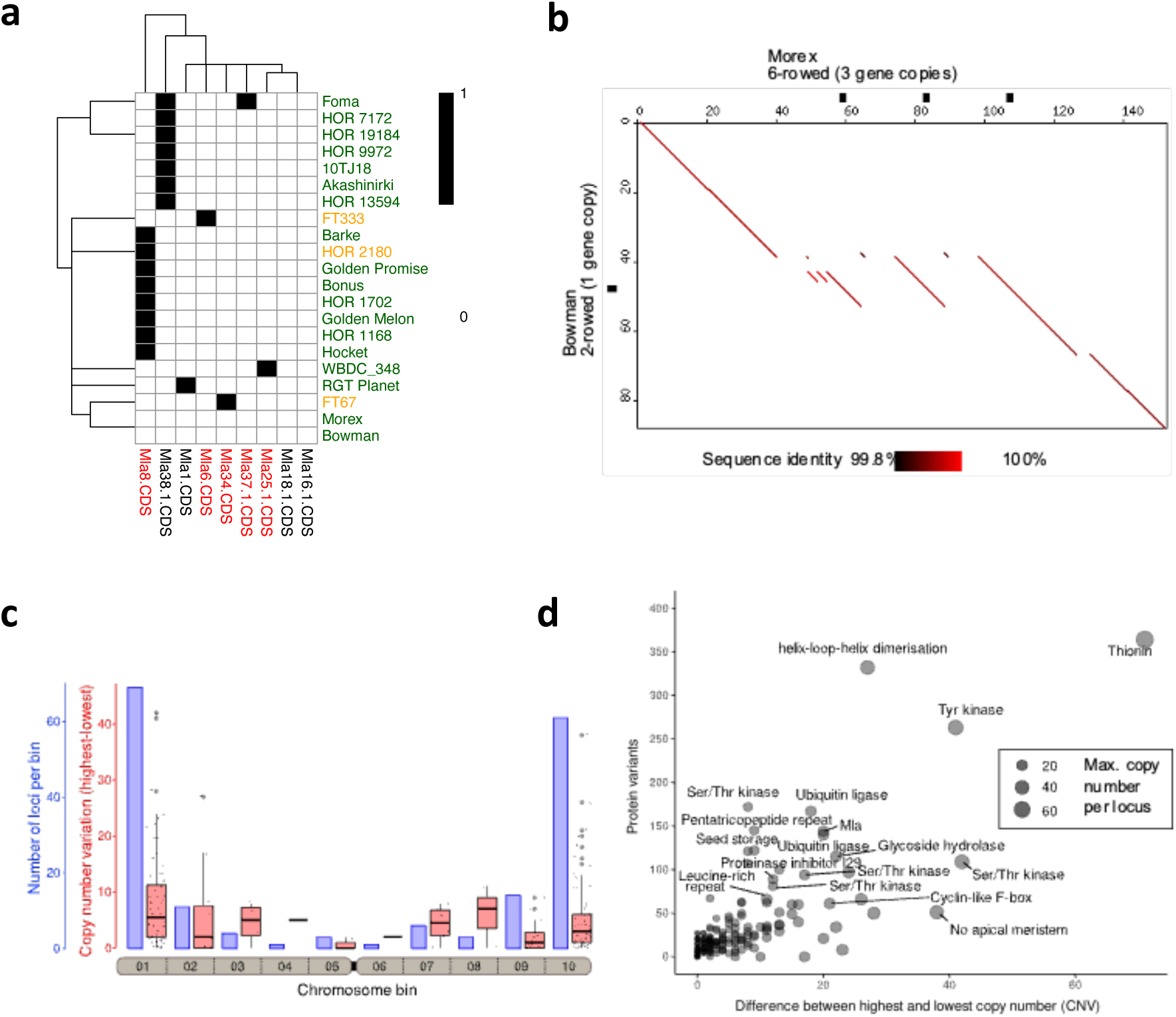
S**t**ructurally **complex loci in the barley pangenome. (a)** Presence/absence of known *Mla* alleles in the barley pangenome. Black and white squares denote presence and absence, respectively. The names of *Mla* alleles (y-axis) and genotypes (x-axis) are coloured according to subfamily and domestication statues, respectively. (green – domesticated; orange – wild). Only the genomes containing known alleles are displayed. **(b)** Dot plot alignment of complex locus Chr04_015772 which contains *Int-c* genes. The plot shows an alignment of Morex (six-rowed barley) and Bowman (two-rowed barley). In Morex, *Int-c* and its surrounding sequence is present in three copies. Genes are indicated as black boxes along the axes of the plot. Individual tandem repeat units are 96-100% identical. **(c)** Complex loci are enriched in distal chromosomal regions. The seven barley chromosomes were divided into ten equally sized bins, and cumulative data for all chromosomes is shown. The bar plot indicates the number of loci, while the box blot shows the extent of CNV for all loci in the bin. **(d)** CNV levels and numbers of encoded protein variants identified in 76 barley accessions. The x-axis shows the level of CNV (i.e. the difference between the accession with the fewest copies to that with the most copies for each locus). The y-axis shows the total number of protein variants identified in all 76 barley accessions. Labels mark genes families with the highest copy numbers or the highest CNV levels.

We employed a gene-agnostic method^24^ to scan the genome sequence of Morex for structurally complex loci harbouring genes, focusing on examples that had evidently caused gene copy number variation across the pangenome via the expansion or collapse of long tandem repeats. A total of 173 loci ranging in size from 20 kb to 2.2 Mb (median: 125 kb) matched our criteria (**Fig. 2c, Supplementary Table 7**). Their copy numbers were variable in the pangenome. The most extreme case was a cluster of genes annotated by homology as thionin genes, which are possibly involved in resistance to herbivory^25^. The locus had as few as three thionin gene copies in the wild barley WBDC103 and up to 78 copies in WBDC199, another wild barley (**Extended Data** Fig. 6d). Genes associated with such complex loci possessed functional annotations suggesting involvement in various biological processes (**Fig. 2c, Supplementary Table 7**). Complex loci were enriched in distal chromosomal regions (**Fig. 2d**). In this regard, they follow the same distal-to-proximal gradient as genetic diversity and recombination frequency. The latter process might play a role in their amplification and contraction owing to unequal homologous recombination between neighboring repeat units^26^ (**Extended Data** Fig. 6a). Molecular dating of the tandem duplications in Morex is consistent with rapid evolution (**Extended Data** Fig. 7): loci with many gene copies appear to have gained them within the last three million years (**Extended Data** Fig. 7c), after the *H. vulgare* lineage split from that of its closest relative *H. bulbosum*^27^. In addition, 63 loci (36.4%) underwent at least one duplication in the last 10,000 years, that is, after domestication (**Extended Data** Fig. 7d). Forty-five loci expanded so recently that the genes they harboured were identical duplicates of each other.

One interesting case of such recent diversification was a duplication at the *HvTB1* locus (also known as *INTERMEDIUM-C* [*INT-C*] or *SIX ROWED SPIKE 5*). HvTB1 is a TEOSINTE BRANCHED 1, CYCLOIDEA, PCF1 (TCP) transcription factor involved in basal branching (tillering) and other aspects of plant architecture in cereal grasses^28–30^. In barley, both tillering and the fertility of lateral spikelets is increased in knock-out mutants^30,31^. Just two alleles, *Int-c.a* and *int-c.b*. dominate in six-rowed and two-rowed forms^30^, respectively, and *HvTB1* is not genetically linked to the *SIX ROWED SPIKE 1* gene. Both alleles of *HvTB1* are thought to be functional and occur also in wild barley^30,32^. These patterns have defied easy explanation. Expression differences owing to regulatory variation have been postulated but not proven^30^. The pangenome adds another twist. *HvTB1* is a single-copy gene in all 22 *H. spontaneum* accessions and 23 two-rowed domesticates except HOR 7385 (**Supplementary Table 8**). Six-rowed forms, however, have up to four copies of a 21 kb segment that contains *HvTB1* and ∼5 kb of its upstream sequence (**Fig. 2b)**. The reference cultivar Morex has three copies, although these were falsely collapsed in previous short-read assemblies of that variety^33^. On top of variable copy numbers, the pangenome revealed six hitherto unknown HvTB1 protein variants (**Extended Data** Fig. 6d**, Supplementary Table 8**). Reduced tillering in maize has been attributed to overexpression of *TB1*. The barley pangenome will help developmental geneticists reveal if copy number gains had analogous effects in six-rowed forms.

## Amplification of α-amylases in malting barley

Among the complex loci we examined, the *amy1_1* locus of α-amylases is arguably the one of greatest economic importance. These enzymes cleave the polysaccharide starch into short-chain forms, which are then digested further into sugars^34^. In both wild and cultivated forms, the speed and efficiency of that process determines the energy supply to and hence the vigor of the young seedling^35^. In grains of domesticated barley, the enzymatic conversion of starch into fermentable sugars by α-amylases initiates the malting process. Barley α-amylases are subdivided into four families, which occupy distinct genomic loci (**Extended Data** Fig. 8a**, Supplementary Tables 9 and 10**). Earlier genome sequences assembled from short reads hinted at the presence of structural variation at the *amy1*_*1* locus on chromosome 6H, respectively, but failed to resolve copy numbers^36^. By contrast, each of our long-read assemblies covered *amy1_1* in a single contig (**Extended Data** Fig. 9a). Copy numbers of *amy1_1* in 76 complete genomes varied between two and eight, with on average more copies in domesticated than in wild forms **(Fig. 3a, b)**. Individual copies were addressable by 21-mers that overlap sequence variants. We counted these 21-mers in the short-reads of 1,315 genotypes and also determined SNP haplotypes around the *amy1_1* locus in these data (**Extended Data** Figs. 8e-f**, 9b**, **Supplementary Tables 11 and 12**). Eight clusters were discernible and could be related to population structure. Three-quarters of hulless barleys were in cluster #7. Six-rowed barleys belonged mostly to clusters #1 and #6. Among 315 European varieties, clusters #5 and #6 were most common. Clusters #3 and #8 with fewer *amy1_1* copies were exclusive to plant genetic resources. Barleys from eastern and central Asian countries tend to have high copy numbers. *Amy1_1* copy numbers were higher on average in elite varieties than in other barleys (**Fig. 3b**). Structural diversity was accompanied by differences in gene sequence owing to SNPs and indels in open reading frames and promoters. The 76 genome assemblies had 94 distinct *amy1_1* haplotypes (**Fig. 3c**, **Extended Data** Fig. 8b, **Supplementary Tables 13-16**). Twelve had insertions of transposable elements (**Supplementary Table 17**). At the protein level, there were 38 unique AMY1_1 isoforms (**Supplementary Tables 18 and 19**), some of which were predicted to affect protein^37^ stability and thereby influence α-amylase activity (**Fig. 3d, e**).

**Figure 3.**
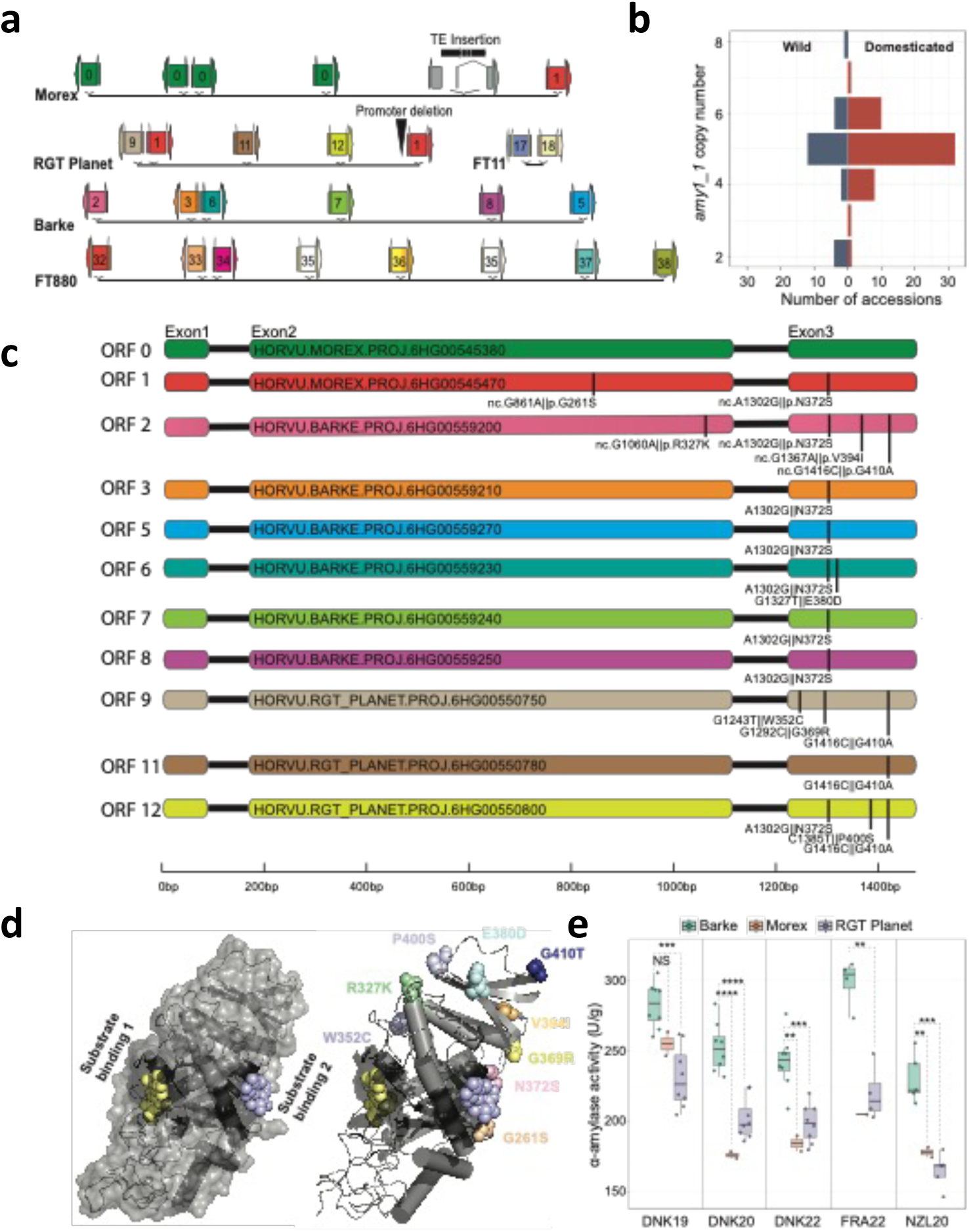
Structural diversity at the *amy1_1* locus and its importance in malting. (a) Simplified structure of the *amy1_1* locus in selected pangenome assemblies. A detailed depiction of the *amy1_1* locus across all 76 assemblies is shown in **Extended Data** Fig. 9a. Identical ORFs have the same colours in **(a)** and **(c)**. **(b)** Distribution of *amy1_1* copy numbers in wild and domesticated accessions of the pangenome. **(c)** Non-synonymous sequence exchanges in 12 non-redundant *amy1_1* ORFs in the malting barleys Morex, Barke and RGT Planet. The positions of sequence variants and respective amino acid variations are marked by black lines. ORF numbers refer to **Supplementary Table 13**. **(d,e**) X-ray crystal structure (pdb: 1BG9; ref. ^36^) of α-amylase bound to acarbose as a substrate analogue (magenta and yellow spheres in panel **(d)**. In panel **(e)**, *amy1_1* amino acid variants (found in Morex, Barke and RGT Planet, **Supplementary Table 20**) are added as coloured spheres. **(f)** α-amylase activity of micro-malted near-isogenic lines (NILs) containing *amy1_1*-Morex, Barke and RGT Planet haplotypes.

We investigated in more detail the elite malting barleys Morex, Barke and RGT Planet (**Fig. 3, Supplementary Tables 20 and 21**). Prior to its use as a genome reference cultivar, Morex was a successful variety in North America. It had six nearly identical (> 99 % similarity) *amy1_1* copies. (**Fig. 3a**). The fifth copy was disrupted by the insertion of a transposable element. Full-length copies were verified by PacBio amplicon sequencing. Barke, a European cultivar, had six full-length copies, albeit of a different haplotype. RGT Planet, currently a successful cultivar in many barley-growing regions around the world, had five copies, one of which was likely to be inactivated by a 32 bp deletion in a pyr-box (CTTT(A/T) core) promoter binding site that is essential for α-amylase transcription^38^. We tested overall α-amylase activity in micro-malting trials with RGT Planet and near-isogenic lines (NILs) that carried Morex and Barke *amy1_1* haplotypes in the genomic background of RGT Planet. It was observed that α-amylase activity was highest in amy1_1-Barke NILs (**Fig. 3e**). The Barke haplotype is common not only in cultivars favored by European maltsters, but also among those from other regions of the world, where barley α-amylases need to be abundant enough to cleave starch from adjuncts such as maize and rice (**Supplementary Table 22**). The patterns of sequence variation at *amy1_1* uncovered by the barley pangenome pave the way for the targeted deployment, possibly even design, of *amy1_1* haplotypes in breeding.

## A regulatory variation controls trichome development

Our last example sits at the intersection of developmental genetics, breeding and domestication. Hairy appendages to grains and awns are conducive to seed dispersal in wild plants, but have lost this function in domesticates^39^. A pertinent example are the hairs on the rachillae of barley grains. In barley, the rachilla is the rudimentary secondary axis of the inflorescence, where multiple grains are set in wheat^40^. In the single-grained spikelets of barley, the rachilla is a thin and hairy thread-like structure nested in the ventral crease of the grains. The long hairs of the rachillae of wild barleys and most cultivated forms are unicellular, while the short hairs of some domesticated types are multicellular and branched (**Fig. 4a, Extended Data** Fig. 10a). This seemingly minor difference in a vestigial organ belies its importance in variety registration trials^41^, where breeders would like to predict the trait with a diagnostic marker. *Short rachilla hair 1* (*srh1*) is also a classical locus in barley genetics^42^. It has been mapped genetically^8,43^ (**Fig. 4b**) and both long-and short-haired genotypes are included in our pangenome. Fine-mapping in a population of 2,398 recombinant inbred lines derived from a cross^36,44^ between cultivars Morex (short, *srh1*) and Barke (long, *Srh1*) delimited the causal variant to a 113 kb interval on the long arm of chromosome 5H (**Fig. 4c, Supplementary Table 23**). Outside of this interval (which is itself devoid of annotated gene models), but within 11 kb of the distal flanking marker is a homolog of a *SIAMESE-RELATED* (*SMR*) gene of the model plant *Arabidopsis thaliana*^45,46^. Members of this family of cyclin-dependent kinase inhibitors control endoreduplication in trichomes of that species. In barley, hair cell development is likewise accompanied by endopolyploidy-dependent cell size increases (**Extended Data** Fig. 10b). The SMR-homolog was expressed in the rachilla’s developing trichomes (**Extended Data** Fig. 10e), but there were no differences between Morex and Barke in the sequence of this otherwise plausible candidate gene. Despite this conflicting evidence, we proceeded with mutational analysis and obtained several mutants using FIND-IT^47^ (**Extended Data** Fig. 10c,d) and Cas9-mediated targeted mutagenesis (**Fig. 4d**, **Supplementary** Fig. 2**, Supplementary Tables 24 and 25**). Mutants of long-haired genotypes with knock-out variants or a nonsynonymous change in a Pro phosphorylation motif (Thr62-Pro63) had short, multicellular rachillae, supporting the idea that the gene in question, *HORVU.MOREX.r3.5HG0492730,* is indeed *HvSRH1*. Sequence variants in *HvSRH1* identified in the pangenome did not lend itself to easy explanation: 18 protein haplotypes caused by 23 non-synonymous variants bore no obvious relation to the phenotype (**Supplementary Table 26**). Thus, we then examined regulatory variation. All 14 short-haired genotypes in the pangenome lacked a 4,273 bp sequence (**Fig. 4c**), which was exceptionally well conserved in long-haired types, with 95% overall identity to Barke. Within this sequence, we found the motif CATCGGATCCTT, matching the sequence [ATC]T[ATC]GGATNC[CT][ATC], which is recognized by regulators of SMR expression in *A. thaliana*^48^. That sequence was repeated five times in Barke. The closest unit in long-haired types was no further than 13.6 kb from the gene, while the minimum distance between the gene and its putative enhancer motif in short-haired types was 22.3 kb, owing to the 4.3 kb deletion (**Fig. 4c**). *HvSRH1* expression during rachilla hair elongation is higher in long-haired than in short-haired genotypes (**Extended Data** Fig. 10f). Gene edits of the enhancer region, guided by the pangenome sequences, will further elucidate the transcriptional regulation of *HvSRH1*.

**Figure 4.**
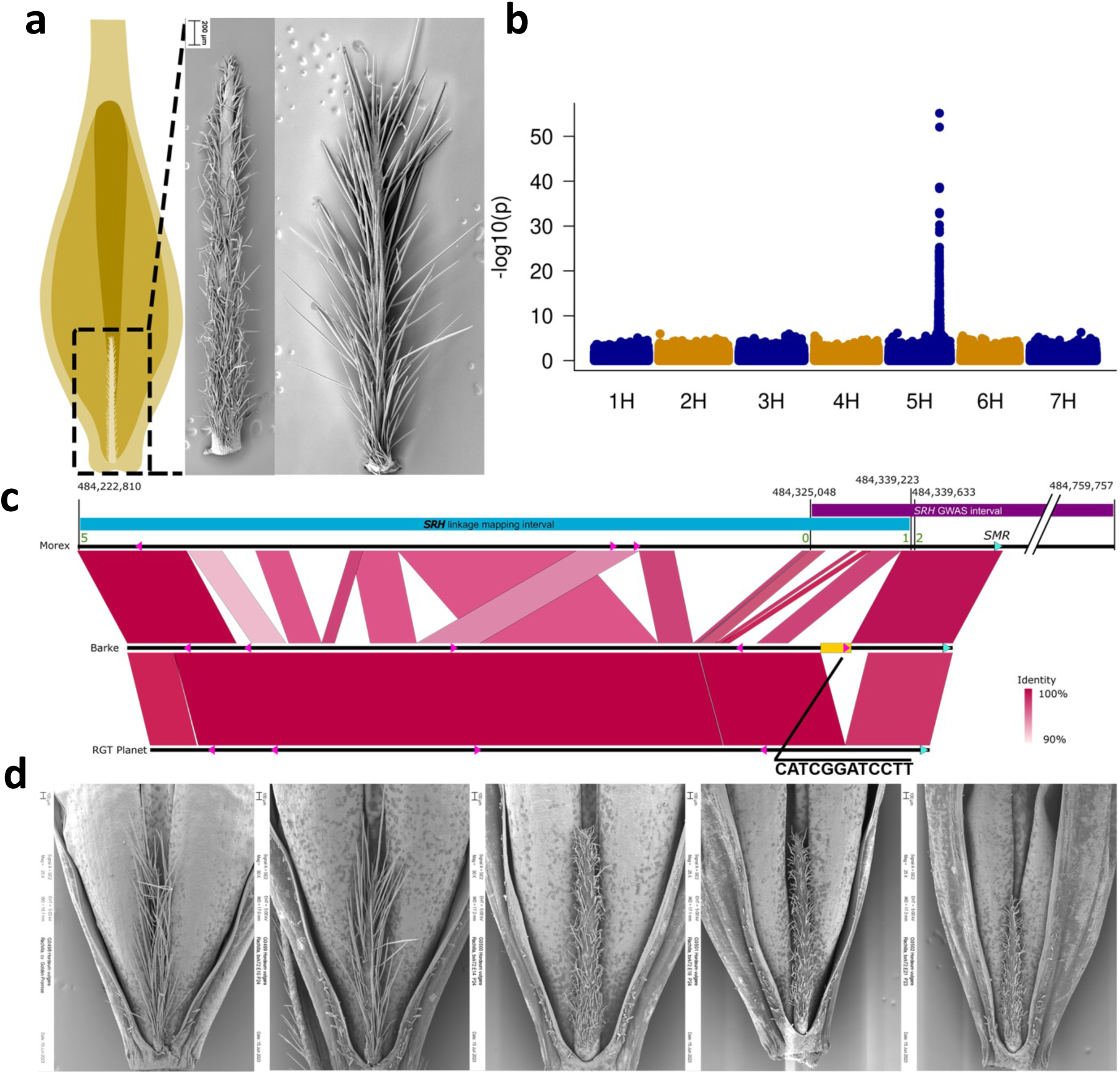
A deletion in an enhancer motif is associated with trichome branching. (a) Schematic drawing of a seed from a hulled and awned barley. The rachilla is a rudimentary structure attached to the base of the seed, representing reduced lateral branches in the barley inflorescence. On the right, scanning electron micrographs are shown of a short-haired and a long-haired rachilla of genotypes Morex and Barke, respectively. **(b)** Genome-wide association study (GWAS) for rachilla hair phenotype in the core1000. **(c)** Top part: schematic representation of the high-resolution genetic linkage analysis at the *Srh1* locus. Blue and purple horizontal bars represent the overlapping biparental and GWAS mapping intervals in reference to the 160 kb physical interval in the Morex genome (black line below the colored bars). Note, the SMR-like gene sits outside the high-resolution biparental mapping interval. Bottom part: connector plot showing orthologous regions in the genotypes Barke (long hairs) and RGT Planet (short hairs). A region harboring a conserved enhancer element (yellow rectangle) is present in Barke, but absent in Morex and RGT Planet. **(d)** Rachilla hair phenotype of the Cas9-induced knock-out mutants of the SMR-like gene. From left to right: wild-type Golden Promise (GP); wild-type segregant from the brhE72P19 family; independent mutant segregants showing the short-hair phenotype.

## Discussion

The recently published human draft pangenome demonstrated how contiguous long-read sequences help make sense of reams of sequence data^49^. Our study on barley pangenome sheds light on crop evolution and breeding. The shortcomings of previous short-read assemblies made it all but impossible to see patterns that now emerge from their long-read counterparts. We were able for the first time to study the evolution of structurally complex loci of nearly identical tandem repeats. Our developmental insights are admittedly still cursory: true to the hypothesis-generating remit of genomics, and at least as many questions were raised as answered. We studied four loci – *Mla*, *HvTB1*, *amy1_1*, *HvSRH1* – and the traits they control: disease resistance, plant architecture, starch mobilization and the hairiness of a rudimentary appendage to the grain. In two of these examples, phenotypic diversity has visibly increased in domesticated forms: there are no six-rowed or short-haired wild barleys. Malting created new selective pressures that only cultivated forms experienced. Novel allelic variation at disease resistance loci is both illustrative of the power of pangenomics and in line with our understanding of how disease resistance genes evolve. Structural variation at *amy1_1* has been known for some time, but previous attempts at resolving the structure of the locus had been thwarted by incomplete genome sequences. Tandem duplications and deletions of regulatory elements, respectively, at *HvTB1* and *HvSRH1* was surprising since for many years barley geneticists considered the loci as monofactorial recessive. Much of the variation seems to have arisen after domestication, either because mutations that appear with clock-like regularity were absent or copy numbers were lower in the wild progenitor than in the domesticated forms. A common concern among crop conservationists is dangerously reduced genetic diversity in cultivated plants^50^. But crop evolution need not be a unidirectional loss of diversity. This study has shown that valuable diversity can arise after domestication. Rapid evolution at structurally complex loci may endow domesticated plants with a means of adaptive diversification that aptly fulfills the needs of farmers and breeders. More diverse crop pangenomes will help us understand how the counteracting forces of past domestication bottlenecks and newly arisen structural variants influence future crop improvement in changing climates.

## Acknowledgments

We are grateful for the technical assistance of Susanne König, Ines Walde, Sabine Sommerfeld, John Fuller, Nicola McCallum and Malcolm Macaulay and thank Thomas Münch, Jens Bauernfeind and Heiko Miehe for IT administration. Andreas Börner supported the development of the core1000 diversity panel. Sequencing of Chikurin Ibaraki 1 was performed at the Institute for Clinical Molecular Biology, Competence Centre for Genomic Analysis (CCGA), Kiel University, Kiel, Germany under the supervision of Dr. Janina Fuß. Sequencing of HOR 4224 was performed at Genomics & Transcriptomics Labor (BMFZ) Heinrich-Heine-Universität, Universitätsklinikum Düsseldorf, Germany under the supervision of Prof. Dr. Karl Köhrer. RGT Planet and Maximus were sequenced at Genomics WA under the supervision of Dr. Alka Saxena and at Novogene, respectively. N.S.,M.M.,K.F.X.M.,M.Spannagl and U.S. were supported by grants from the German Ministry of Research and Education (BMBF, 031B0190 and 031B0884). D.P. was supported by BMBF grant 031B0199B and the German Federal Ministry of Food and Agriculture (grant 2818BIJP01). U.S.’s research is supported by the German Research Foundation (DFG, grant 442032008). C.D., Q.L., P.R.P. and B.S. gratefully acknowledge support from the Carlsberg Foundation to B.S. (grants CF15-0236, CF15-0476, CF15-0672) and thank F. Lok, S. Knudsen, G. B. Fincher and K. G. Jørgensen for providing valuable scientific thoughts and discussions on barley α-amylases and malt quality. R.W., M.B., M.Schreiber, W.G., R.Z. and C.S. received funding from the Rural and Environment Science and Analytical Services Division (RESAS, grant KJHI-B1-2). W.G. and R.Z. were supported by grant BB/S020160/1 from the Biotechnology and Biological Sciences Research Council (BBSRC). C.P. acknowledges support from Genome Canada and Genome Prairie. B.K. acknowledges a grant from the Swiss National Science Foundation (310030_204165). C.L., K.C. and P.L. were supported by a grant from the Grain Research and Development Corporation (UMU1806-002RTX), by the Department of Primary Industry and Regional Development Western Australia and by Pawsey Australia (for computational resources). K.P. and T.S. received a grant from DFG (HAPPAN, 433162815). G.S.B. has received support from the Saskatchewan Ministry of Agriculture (grant ADF20200165) and from the Saskatchewan Barley Development Commission, Western Grains Research Foundation, Alberta Barley Commission and the Manitoba Crop Alliance (grant ADF20210677). A.B. and W.B. are recipients of the TUGBOAT grant from the Agriculture and Agri-Food Canada – Genomics Research and Development Initiative and the grant “Unlocking barley endophyte microbiome to enhance plant health and grain quality” from Agriculture and Agri-Food Canada – A-Base – Foundational Science. M.H. is supported by the Swedish Research Council (VR 2022-03858), the Swedish Research Council for Environment, Agricultural Sciences, and Spatial Planning (FORMAS 2018-01026), the Erik Philip-Sörensen Foundation and the Royal Physiographic Society in Lund. K.Sato’s research is funded by the Japan Science and Technology Agency (grant no. 18076896) and the Japan Society for the Promotion of Science (JSPS, grant no. 23H00333). K.Shirasawa is supported by the JSPS grants nos. 22H05172 and 22H05181. S.S. acknowledges the Research Support Project for the Next Generation at Tottori University. B.S. is supported by the Lieberman-Okinow Endowment at the University of Minnesota and S.G.K. by baseline funding at KAUST. The authors acknowledge the Research/Scientific Computing teams at The James Hutton Institute and NIAB for providing computational resources and technical support for the “UK’s Crop Diversity Bioinformatics HPC” (BBSRC grant BB/S019669/1), use of which has contributed to the results reported in this paper.

## Author contributions

N.S. and M.Mascher designed the study. N.S. coordinated experiments and sequencing. M.Mascher and M.J. supervised sequence assembly. M.Spannagl and K.F.X.M. supervised annotation. U.S. supervised data management and submission. Selection of genotypes: A.B., W.B., G.S.B., K.J.C., Y.G., M.H., B.K., S.G.K., P.L., C.L., M.Mascher, A.M., G.J.M., D.P., K.P., C.J.P., S.S., K.Sato, T.S., B.S., N.S., R.W. Genome sequencing: B.B., A.H., S.I., M.K., C.L., S.P., S.S., K.Sato, T.S., M.Schreiber, K.Shirasawa, N.S., S.W., X.Z. Sequence assembly: B.C., H.H., M.J., G.K.-G., M.Mascher, S.P., K.Sato, T.S., J.F.T. Transcriptome sequencing and analysis: W.G., A.H., S.P., C.S., N.S., R.Z. Annotation: H.G., G.H., N.K., T.L., K.F.X.M., M.Spannagl. Analysis and interpretation of structural variants: M.B., B.C., J.-W.F., Y.G., M.J., C.L., M.P.M., A.M., S.P., H.P., K.P., T.S., M.Schreiber, P.W. Analysis of complex loci: B.J., B.K., M.T.R.-W., N.S., T.W. Analysis of *amy1_1* locus: K.B., C.D., M.E.J., B.J., M.J., S.M.K., Q.L., M.P.M., E.M., P.A.P., P.R.P., B.S., H.C.T., M.T.S.N., D.V., C.V., M.W.R. *srh1* analysis: C.D., M.E.J., I.H., R.E.H., M.J., R.K., J.K., Q.L., M.Melzer, H.P., L.R., P.R.P., T.R., B.S., N.S., H.C.T., C.T., C.V., M.W.R. Data management and submission: M.B., M.F., A.F., M.J., P.K., M.Mascher, D.S., U.S. Writing: M.B., C.D., M.E.J., M.J., Q.L., M.Mascher, N.S., T.W. Coordination: C.D., M.Mascher, N.S. All authors read and commented on the manuscript.

## Competing interests

K.B., C.D., M.E.J., S.M.K., Q.L., E.M., P.R.P., B.S., H.C.T., M.T.S.N., C.V., M.W.R. are current or previous Carlsberg A/S employees. P.A.P. and D.V. are SECOBRA Recherches employees. All other authors declare no competing interests.

## List of supplementary items

**Supplementary Figure 1:** Structure and copy number variation at *Mla* at different thresholds for alignment similarity.

**Supplementary Figure 2:** Targeted mutagenesis at *HvSRH1*.

**Supplementary Table 1:** Passport data of 76 genotypes and statistics and accession codes of their long-read assemblies

**Supplementary Table 2:** Accession codes of transcriptome data

**Supplementary Table 3:** Gene annotation statistics

**Supplementary Table 4:** Gene ontology enrichment in genepool-specific orthologous groups

**Supplementary Table 5**: Passport data of 1,315 genotypes sequenced with short reads, accession codes and mapping stats

**Supplementary Table 6**: Statistics and accession codes of 46 gene-space assemblies

**Supplementary Table 7:** List of 173 structurally complex loci.

**Supplementary Table 8:** Allelic profiles of 76 barley accessions at the 4H_015772 locus (*Int-c*) and at *Vrs1*.

**Supplementary Table 9:** α-amylase gene IDs and chromosomal locations in Morex.

**Supplementary Table 10:** Sequence identity matrix of germination-related *amy1*, *amy2* and *amy3* genes in the Morex genome. *amy4* genes involved in general starch metabolism were excluded due to low sequence identity with other α-amylases.

**Supplementary Table 11:** SNP haplotype clustering analyses and k-mer based *amy1_1* copy number estimation in 1,000 plant genetic resources

**Supplementary Table 12:** SNP haplotype clustering analyses and k-mer based *amy1_1* copy number estimation in 315 European elite cultivars

**Supplementary Table 13:** Overview of *amy1_1* unique ORFs (start to stop codon including intron). HORVU.MOREX.PROJ.6HG00545380 was used as the reference.

**Supplementary Table 14:** Overview of *amy1_1* ORF haplotypes (ORFHap#).

**Supplementary Table 15:** Overview of *amy1_1* unique CDS. HORVU.MOREX.PROJ.6HG00545380 was used as the reference.

**Supplementary Table 16:** Overview of *amy1_1* CDS haplotypes (CDSHap#).

**Supplementary Table 17:** *amy1_1* genes with insertions of transposable elements in the genome assemblies.

**Supplementary Table 18:** Overview of *amy1_1* unique proteins. HORVU.MOREX.PROJ.6HG00545380 was used as the reference.

**Supplementary Table 19:** Overview of *amy1_1* protein haplotypes (ProtHap#).

**Supplementary Table 20:** Amino acid variation in three *amy1_1* haplotypes commonly found in elite varieties (Morex, Barke and RGT Planet).

**Supplementary Table 21:** DynaMut2 prediction of protein stability changes by amino acid variants found in three BPGv2 representatives of widely used *amy1_1* haplotypes found in elite breeding material (Morex, Barke and RGT Planet).

**Supplementary Table 22:** *amy1_1*-Barke haplotype genotyping of AMBA(American Malting Barley Association)-recommended two-row spring malting barley varieties accredited for adjunct brewing.

**Supplementary Table 23:** PACE markers designed in this study.

**Supplementary Table 24:** Oligonucleotides used for gRNA cloning and for PCR amplification of the target region.

**Supplementary Table 25:** Summary of lesions induced in *HvSRH1* by Cas9-mediated targeted mutagenesis.

**Supplementary Table 26:** *srh1* phenotypes and *HvSRH1* gene coordinates in 76 pangenome accessions.

## Extended Data Figures

**Extended Data Figure 1:**
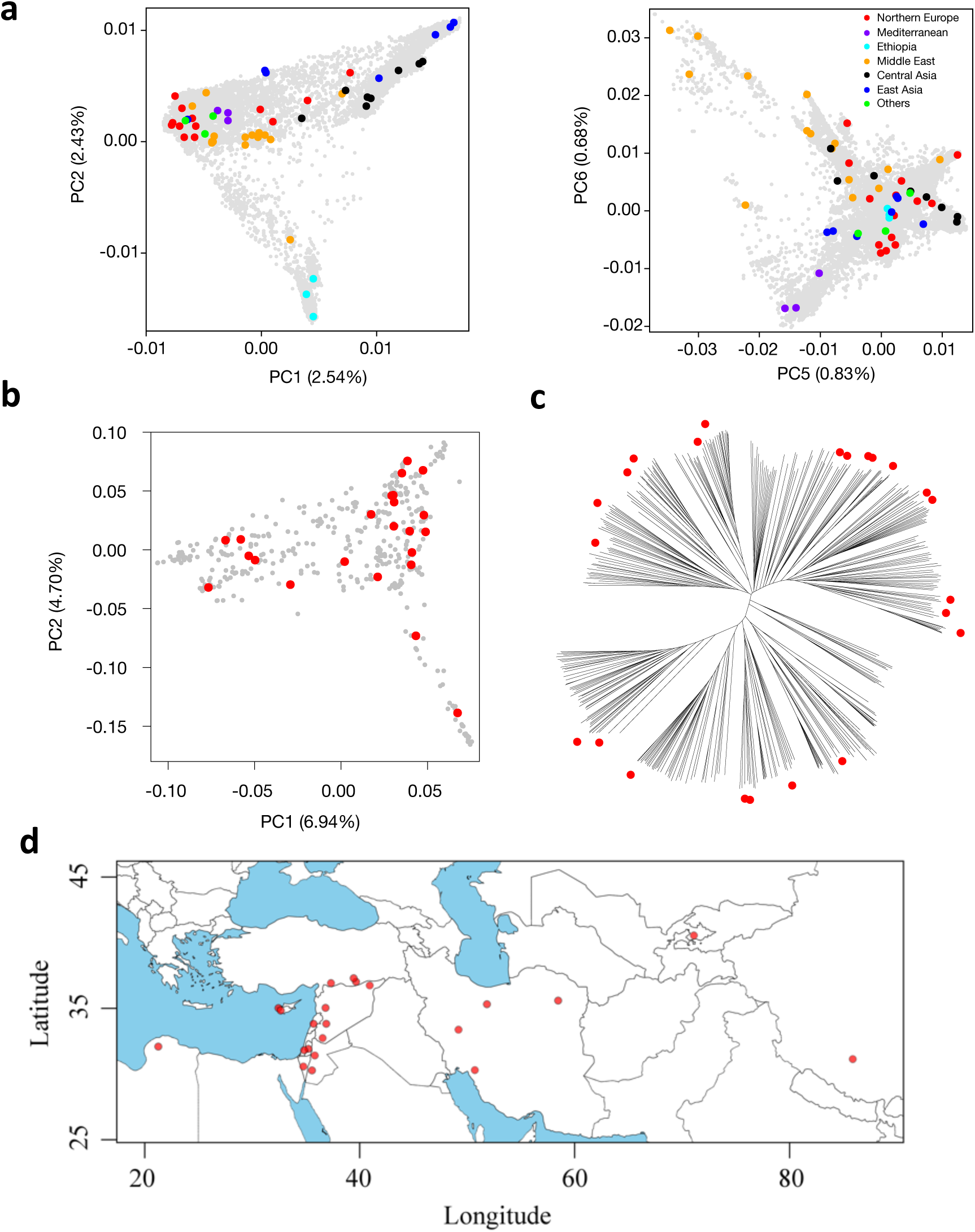
A globally representative diversity panel of domesticated and wild barley. (a) Higher principal components (PC) of the barley diversity space with pangenome accessions highlighted. **(b)** The first two PCs of the diversity space of 412 wild barley (*Hordeum vulgare* subsp. s*pontaneum*) with pangenome accessions highlighted. **(c)** Neighbor-joining phylogenetic tree of those wild barleys. The branch tips corresponding to accessions selected for the pangnome are marked with red circles. The proportion of variance explained by each PC in panels **(a)** and **(b)** is given in the axis labels. **(d)** Map showing the collection sites of wild accessions (n=23) included in the pangenome panel.

**Extended Data Figure 2:**
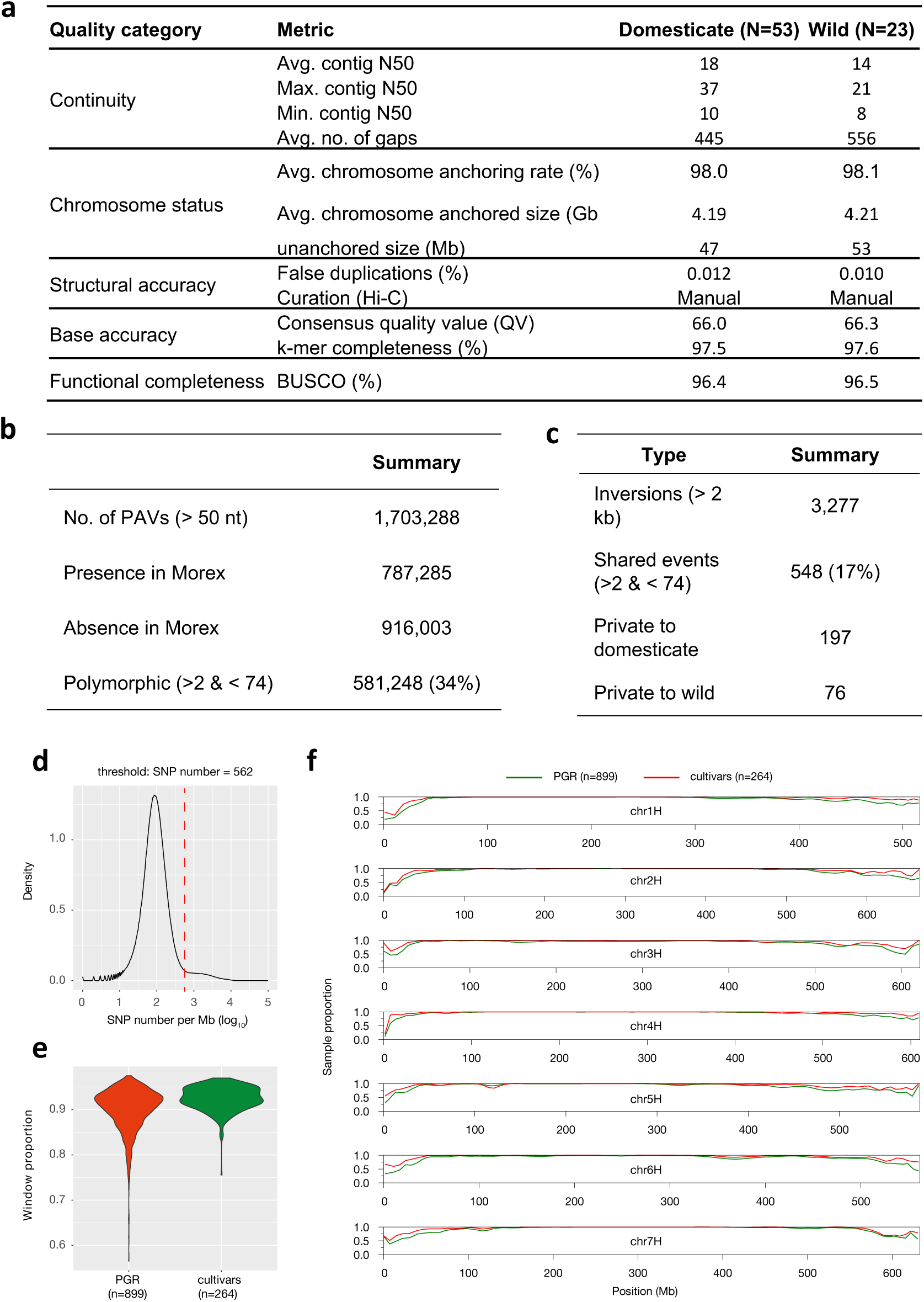
A pangenomic diversity map of barley. (a) Assembly statistics of 76 chromosome-scale reference genomes sequences. **(b)** Counts of presence/absence variants. **(c)** Counts of inversion polymorphisms spanning 2 kb or more. **(d)** Selection of threshold based on pairwise differences (number of SNPs per Mb) for the binary classification into similar/dissimilar haplotypes. **(f)** The proportion of samples with a close match to one of the 76 pangenome accessions is shown for plant genetic resources (PGR) and elite cultivars in sliding windows along the genome (size: 1 Mb, shift: 500 kb). **(h)** Distribution of the share of similar windows in individual PGR and cultivar genomes.

**Extended Data Figure 3:**
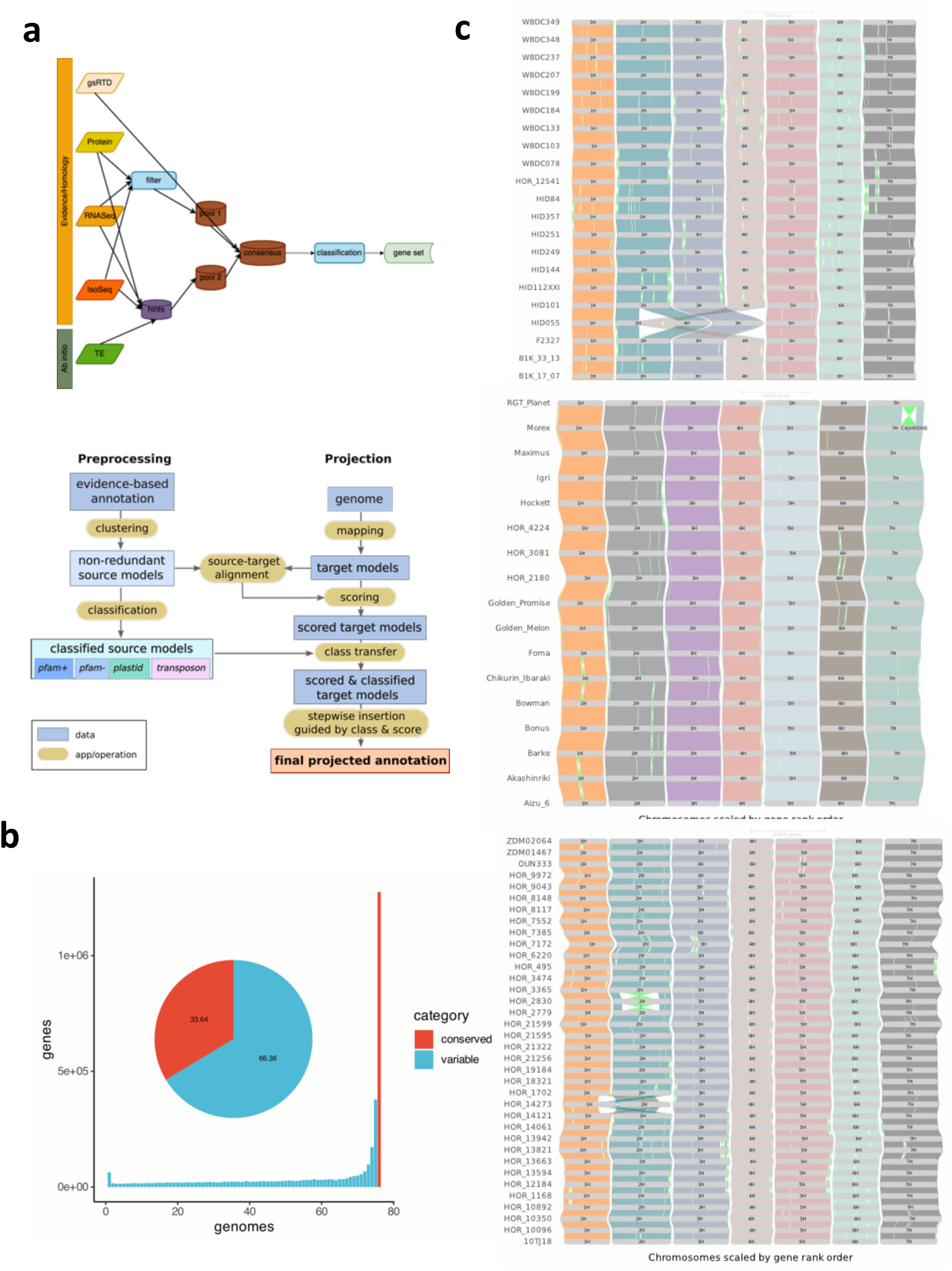
Gene annotation and orthologous framework. (a) Workflow for annotating, projecting and clustering gene models. The upper panel describes the workflow for the de-novo gene predictions, the lower panel for the gene projections **(b)** Histogram showing the number of pagenome genotypes contributing to individual hierarchical orthologous groups (HOGs). The pie chart shows the ratio between conserved and variable genes. **(c)** GENESPACE alignments of 76 barley genomes, grouped by wild barley, cultivated barley and landraces.

**Extended Data Figure 4:**
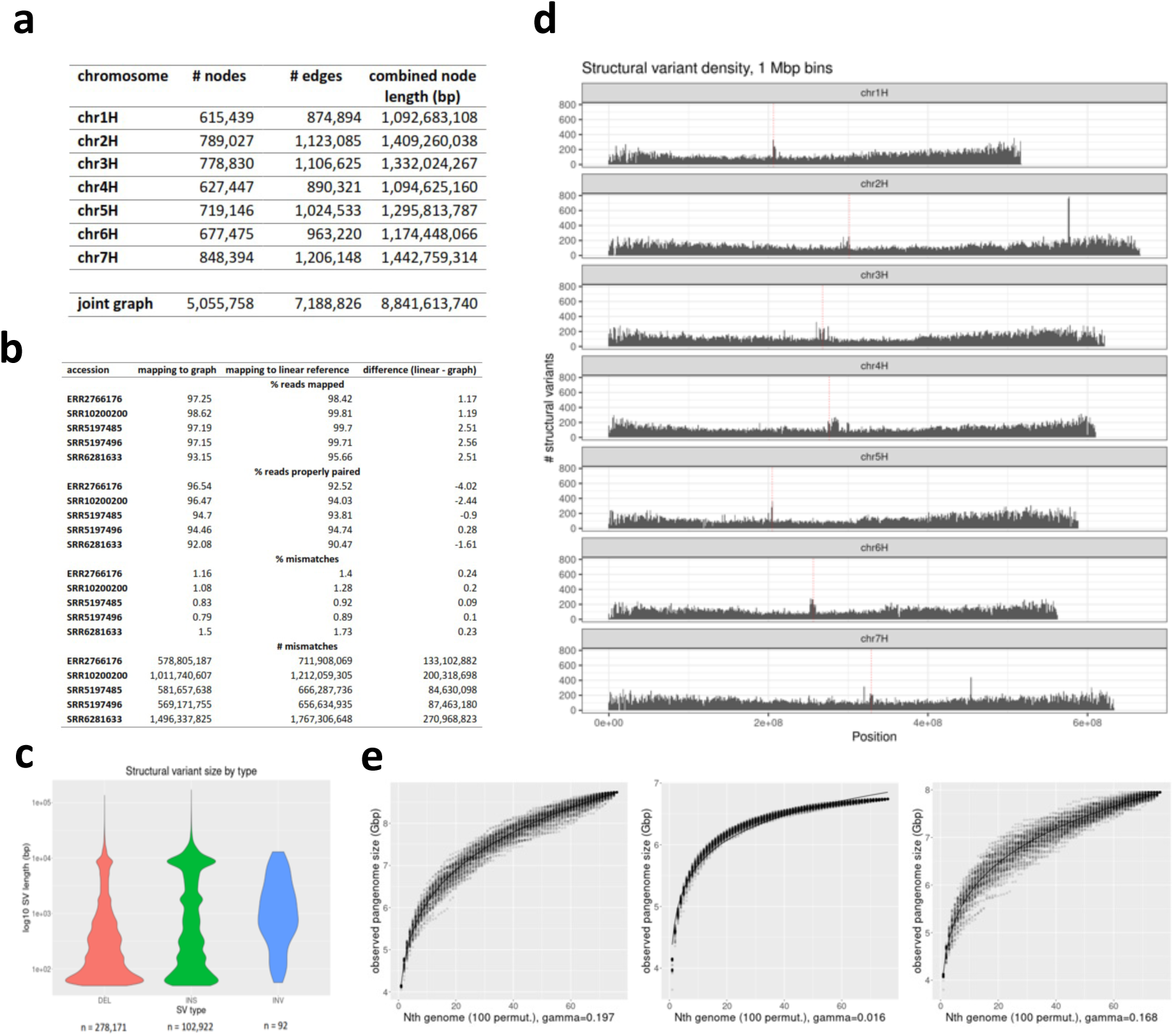
Graph-based pan-genome analysis with Minigraph. **(a)** Descriptive statistics per chromosome and for joint graph. **(b)** Comparative statistics of read mappings from five publicly available Illumina whole genome shotgun sequence read runs against the pan-genome graph and the MorexV3 linear reference sequence. **(c)** Size distribution of structural variants (SVs) in graph. **(d)** Chromosomal distribution of SVs. Centromere positions are indicated by vertical dashed lines in red. **(e)** Pan-genome graph growth curves generated with the odgi heaps tool. One hundred permutations were computed for each number of genomes included. Values of gamma > 0 in Heaps’ law indicate an open pan-genome. Plots shown are for all accessions (left, n = 76), domesticated accessions only (cultivars + landraces, centre, n = 53) and *H. spontaneum* accessions (right, n = 23).

**Extended Data Figure 5:**
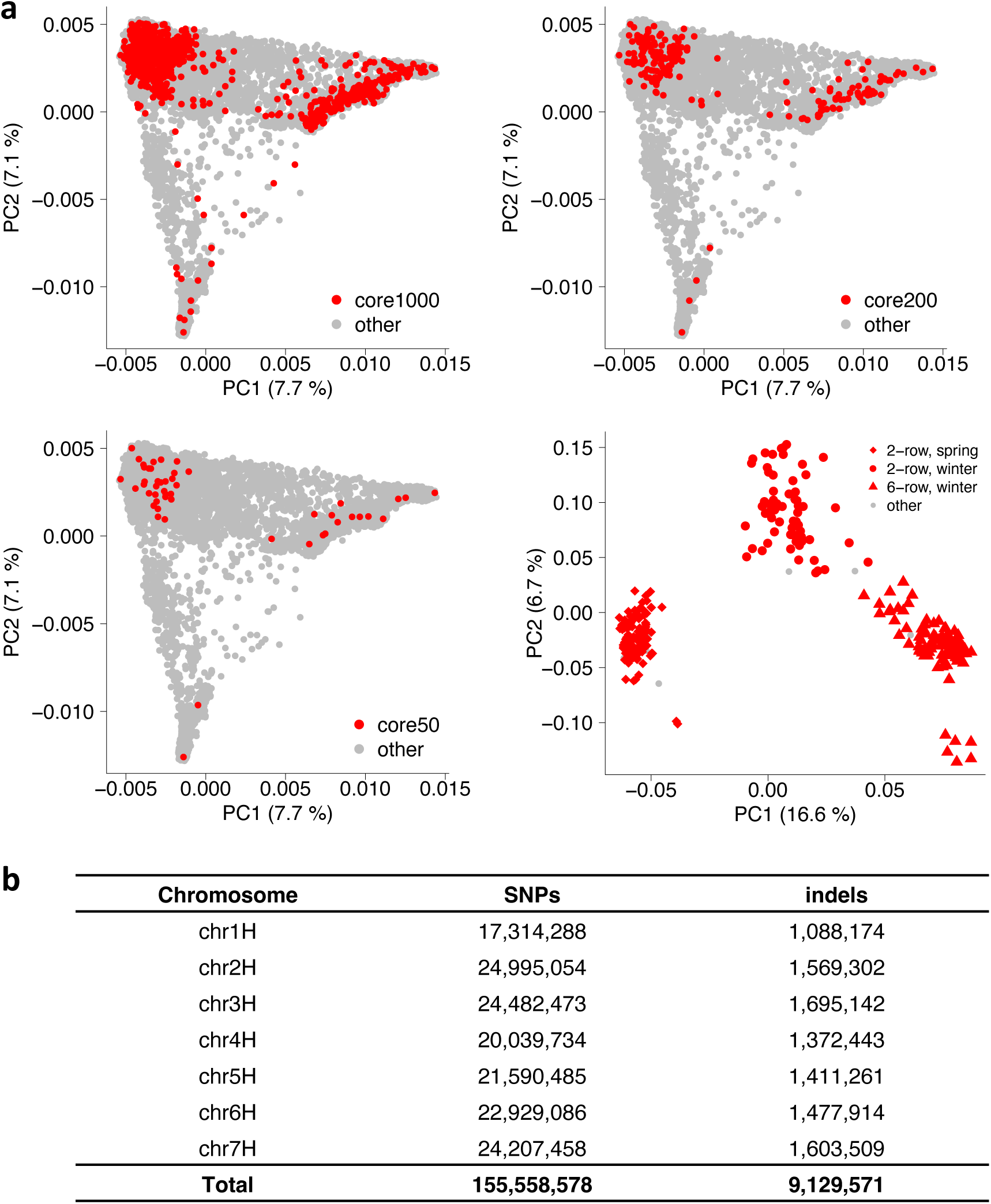
Short-read data complement the pangenome infrastructure. (a) Accessions selected for short-read sequencing. Nested coresets of 1000, 200 and 50 accessions (core1000, core200, core50) are shown in the global diversity space of barley as represented by a principal component (PCA). The top-right subpanel shows a PCA of 315 elite cultivars. Accessions are according to genepool (2-rowed spring, 2-rowed winter, 6-rowed winter). The proportion of variance explained by the PCA is shown in the axis labels. **(b)** Counts of single-nucleotide polymorphisms (SNPs) and short insertions and deletions (indels) detected in those data.

**Extended Data Figure 6.**
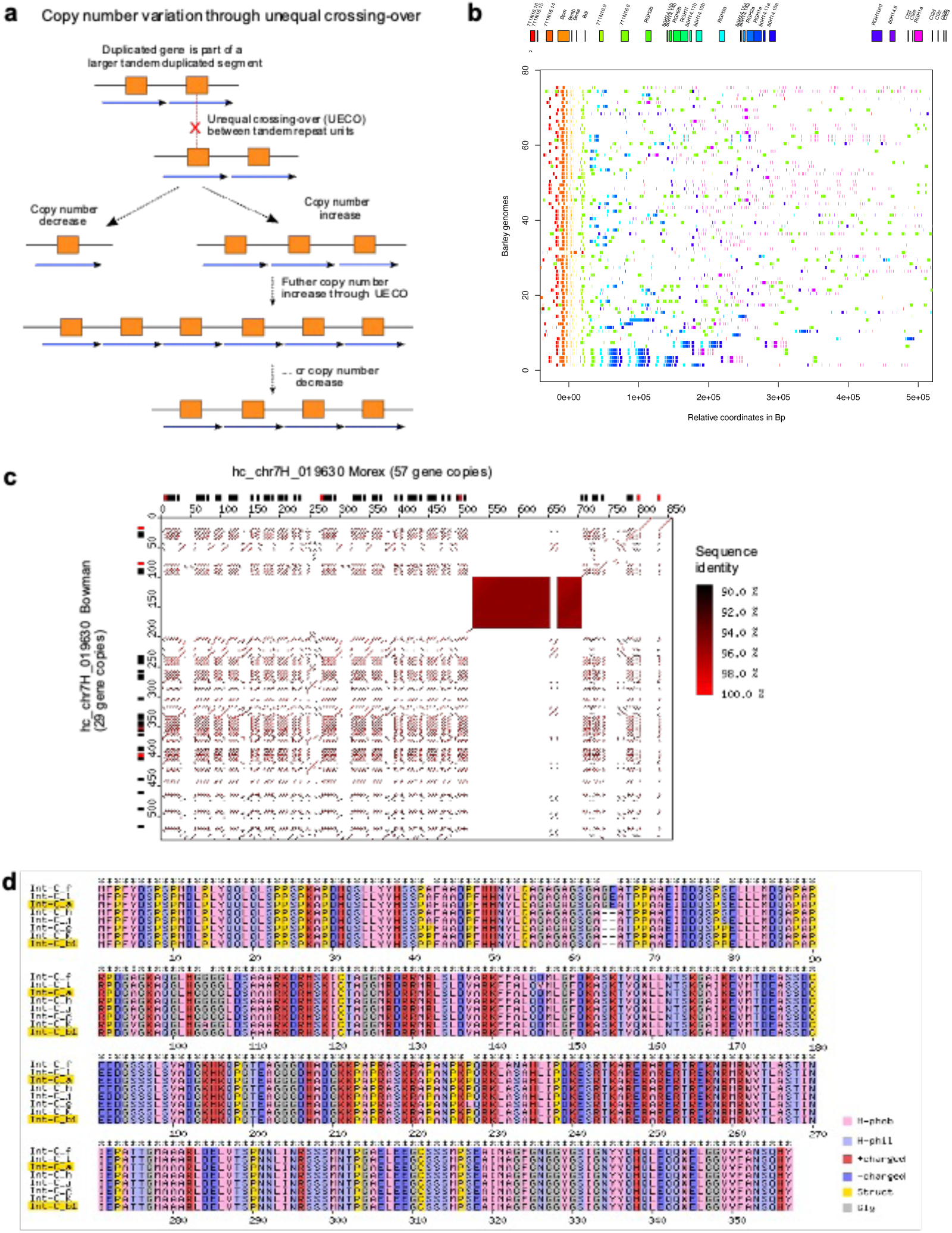
Complex loci are hot spots for copy number variation (CNV). (a) The schematic model shows how, once an initial duplication is established, unequal homologous recombination (unequal crossing-over, UECO) between repeat units can lead to rapid expansion and contraction of the loci, thereby leading to CNV of genes. **(b)** Structure of the *Mla* region across the 76 pangenome accessions. The gene models present in the Morex genome are shown on top. **(c)** Dot plot alignment of the example locus chr7H_019630 which contains a cluster of thionin genes. The sequences of cv. Morex (horizontal) and wild barley HID101 (vertical) were aligned. Predicted intact genes are indicated as black boxes along the left and top axes. Predicted pseudogenes are shown in red. The axis scale is kb. The filled rectangle at positions ∼520-720 kb in Morex represents an array of short tandem repeats which does not contain genes and does not have sequence homology to the gene-containing tandem repeats of the locus. **(d)**. Predicted protein variants of *Int-c* (*HvTB1*) genes. Previously described alleles are highlighted in yellow. Color code: H-phob: Hydrophopic aa, H-phil: Hydrophilic aa, +charged: positively charged aa, – charged: negatively charged aa, Struct: structural aa, Cystein or Prolin, Gly: Gycin.

**Extended Data Figure 7.**
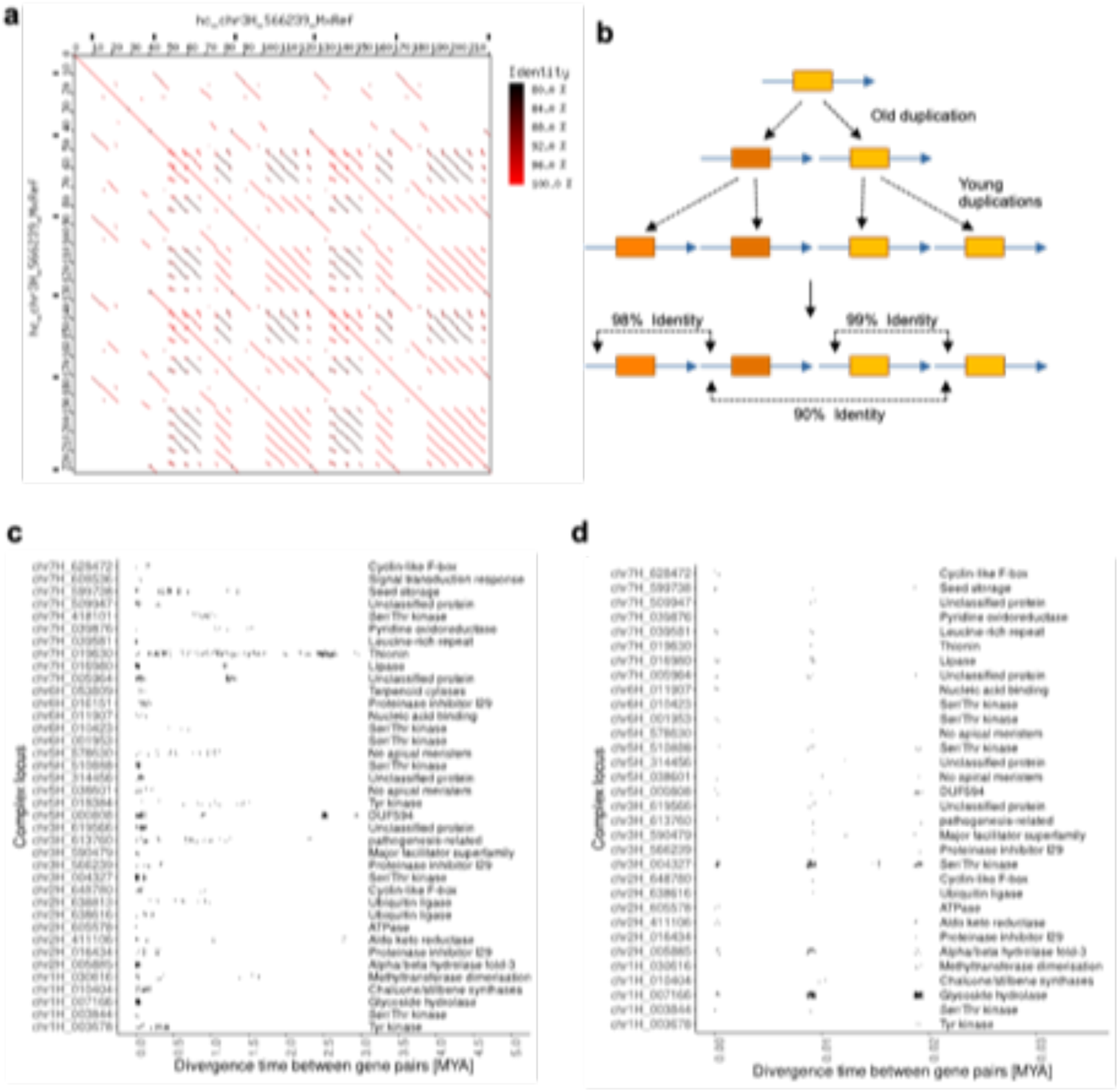
Molecular dating of divergence times between duplicated gene copies in complex loci. **(a)** Dot plot example of locus hc_chr3H_566239 which underwent multiple waves of tandem duplications, which is reflected in varying levels of sequence identity between tandem repeats (color-coded). **(b)** Schematic mechanism for how different levels of sequence identity between tandem repeats evolve. In the example, an ancestral duplication was followed by two independent subsequent duplications, leading to varying levels of sequence identity between tandem repeat units. Genes are indicated as orange boxes while blue arrows indicate the tandem repeats they are embedded in. **(c)** Divergence time estimates between duplicates gene copies in complex loci. Shown are only those complex loci which have at least six tandem-duplicated genes. Each dot represents one divergence time estimate for a duplicated gene pair from the respective locus. The x-axis shows the estimated divergence time in million years. At the right-hand side, classification of proteins encoded by genes in the locus are shown. Note that several loci had multiple waves of gene duplications over the past 3 million years. **(d)** Subset of those loci shown in **(c)** that had at least one gene duplication within the past 20,000 years. The divergence time estimates appear in groups, since they represent the presence of 0, 1 and 2 nucleotide substitutions, respectively, in the approx. 4 kb of aligned sequences that were used for molecular dating.

**Extended Data Figure 8.**
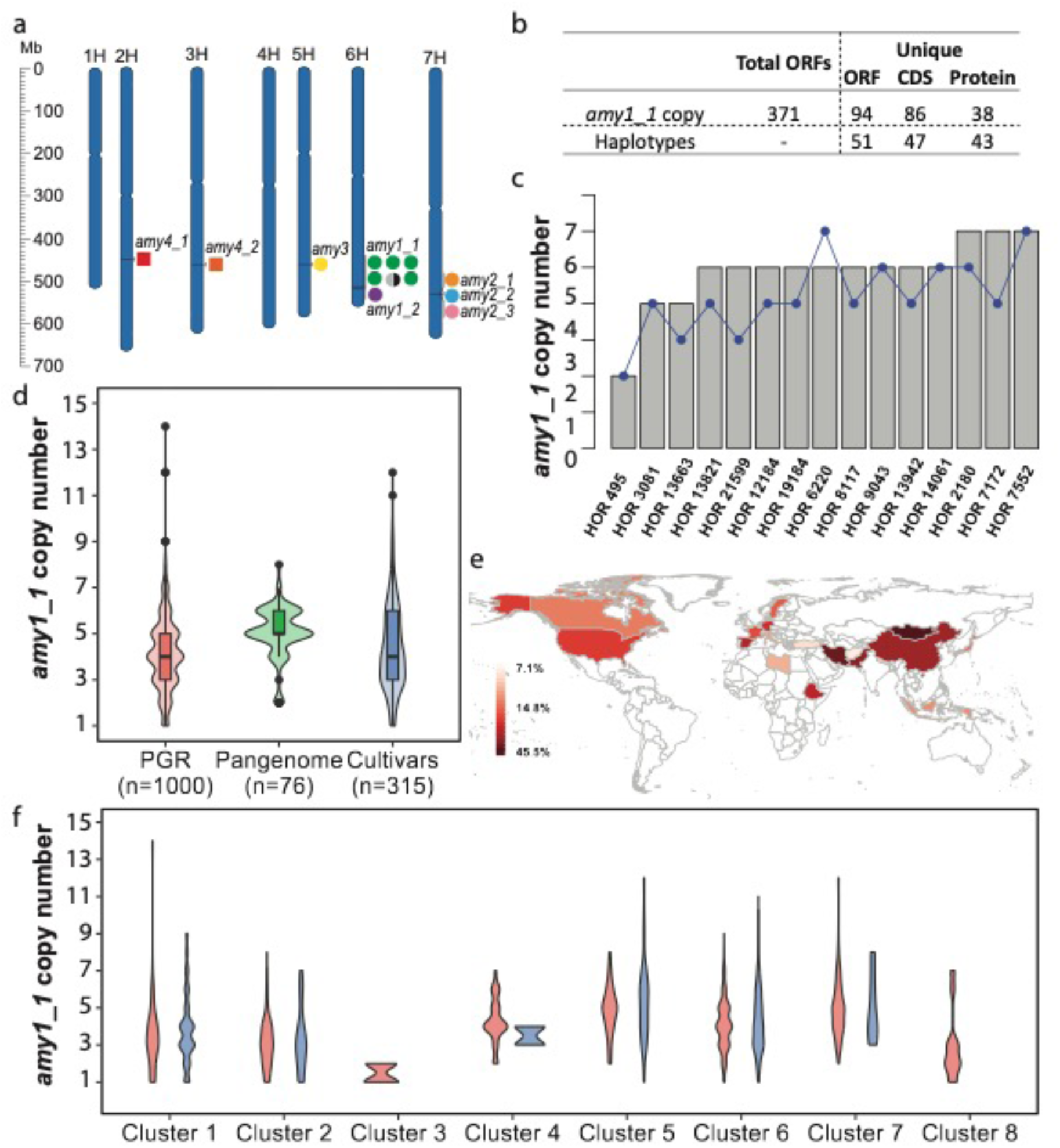
*amy1_1* locus structure and copy number in 76 assemblies and 1,315 whole genome sequenced accessions. (a) Chromosomal location of 12 α-amylase genes in the MorexV3 genome assembly. **(b)** Summary of *amy1_1* locus sequence diversity in 76 pangenome assemblies (**Supplementary Tables 13-16, 18-19**). Total *amy1_1* ORFs in pangenome and unique copies and haplotypes of ORF, CDS and protein. Haplotype denotes unique combinations of ORF, CDS and protein in individual accessions. **(c)** Comparison of *amy1_1* copy numbers identified in the pangenome assemblies versus *k*-mer based estimation from raw reads. Grey bars denote copy number from pangenome, blue dots denote *k*-mer estimated copy number. **(d)** *amy1_1* copy number estimation in 76 pangenome assemblies (“Pangenome”), 1,000 whole-genome sequenced plant genetic resources (“PGR”), and 315 whole-genome sequenced European elite cultivars (“Cultivars”) using *k*-mer based methods. **(e)** Distribution of accessions with *amy1_1* copy numbers >5 per country (as percentage of total accessions in country for countries with ≥10 accessions). **(f)** *amy1_1* copy number within each haplotype cluster (see E**xtended Data** Figure 9b). Red color refers to 1,000 plant genetic resource accessions, green refers to 76 pangenome accessions and blue refers to 315 European elite cultivars. Cluster #5, #6 and #7 contain Barke, RGT Planet and Morex, respectively.

**Extended Data Figure 9.**
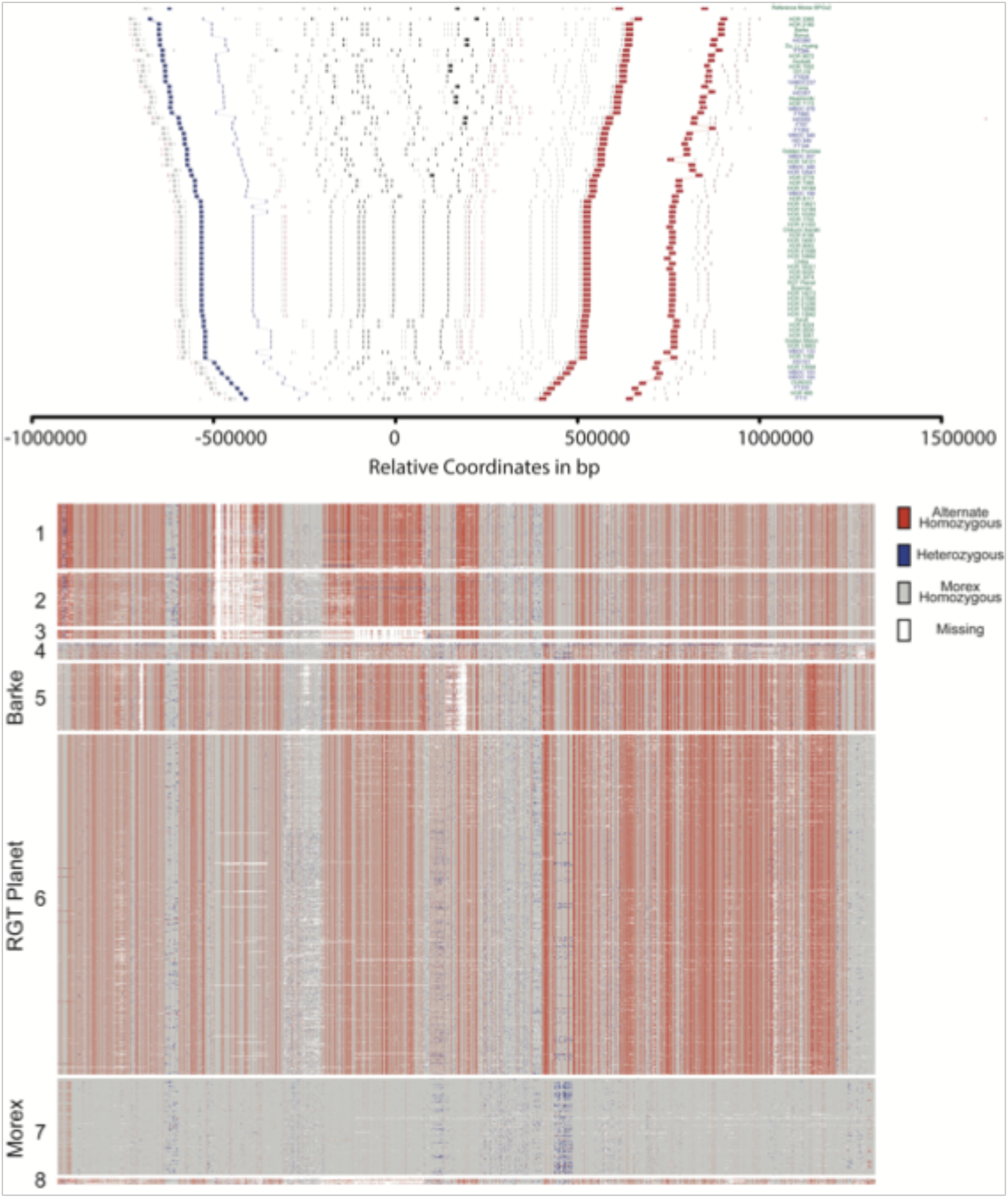
Haplotype structure of the *amy1_1* locus. (a) Structural diversity in the vicinity of *amy1_1* in the 76 pangenome assemblies. Each line shows the gene order in the sequence assembly of one genotype. The Morex V3 reference is shown on top. Coloured rectangles stand for gene models extracted from BLAST alignments against the corresponding gene models in MorexV3. Black rectangles represent *amy1_1* homologs and grey rectangles other genes. Blue and red rectangles represent marker genes used to define the synteny, delimit the region, and sort the accessions based on the distance between endpoints. Lines connect genes models between different genomes. Accession names are given on the right axis and are coloured according to type (blue – wild, green – domesticated). In HOR 8148, five copies assigned to 6H are shown. Two copies assigned to an unanchored contig are not shown. **(b)** SNP haplotype clusters at the *amy1_1* locus among 1,315 genomes of domesticated and wild barley accessions, including genomes of 315 elite barley cultivars. The 6H:516,385,490-517,116,415 bp in the Morex V3 genome sequence is shown. Haplotype clusters #5, #6 and #7 contain the elite malting varieties Barke, RGT Planet and Morex, respectively.

**Extended Data Figure 10:**
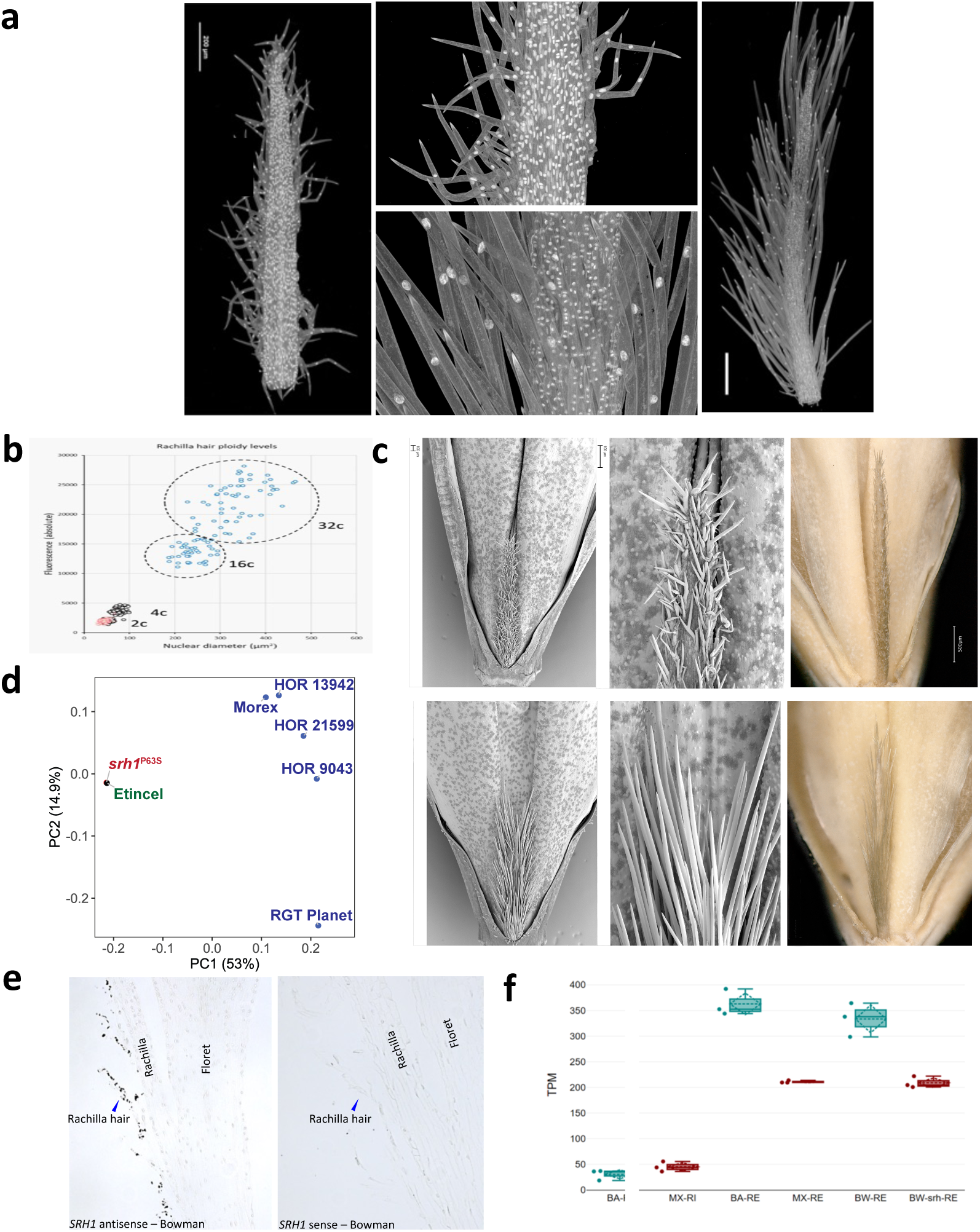
Genetic dissection of the *srh1* locus. (a) Light microscopy of short-and long-haired rachillae at developmental stage W8.5-9 using DAPI staining to visualize the nuclei. Size differences of nuclei in epidermal and trichome cells are very obvious. **(b)** Densitometric measurement of DNA content in epidermal and trichome cells of DAPI stained rachillae of genotypes Morex and Barke, respectively. While trichome cells in short-haired rachillae undergo only one cycle of endoreduplication, the cells in long haired trichomes show eight to sixteen-fold higher DNA contents than epidermal cells indicating three to four cycles of endoreduplication. **(c)** *srh1* mutant discovery. FIND-IT screenings identified a mutant with short-fuzzy hairs (top) in the background of the long-haired variety Etincel (bottom). The mutants are a P64S non-synonymous sequence exchange. Scale bar – 1mm. **(d)** Principal coordinate analysis of SNP array genotyping data of different barley genotypes. Etincel and its mutant cluster together, proving their isogenicity. **(e)** mRNA *in situ* hybridization of *HvSRH1* in longitudinal spikelet sections of Bowman with anti-sense (left) and sense (right) probes. The blue arrow indicates the position of a rachilla hair. **(f)** *HvSRH1* transcript abundance in RNA sequencing data of rachilla tissue in Barke (BA, long-haired), Morex (MX, short-haired), Bowman (BW, long-haired) and a short-haired near-isogenic line of Bowman (BW-srh). Samples were taken at two development stages: rachilla hair initiation (RI) and elongation (RE). Abundance was measured as transcripts per million (TPM).

## Plant growth and high molecular weight DNA isolation

Twenty-five seeds each from the selected accessions (**Supplementary Tables 1 and 6**) were sown on 16 cm diameter pots with compost soil. Plants were grown under greenhouse conditions with sodium halogen artificial 21°C in the day for 16 hrs and 18°C at night for 8 hrs. Leaves (8 g) were collected from 7-day old seedlings, ground with liquid nitrogen to a fine powder and stored at –80°C.

High molecular weight (HMW) DNA was purified from the powder, essentially as described^1^. Briefly, nuclei were isolated, digested with proteinase K and lysed with SDS. Here, a standard watercolor brush with synthetic hair (size 8) was used to re-suspend the nuclei for digestion and lysis. HMW DNA was purified using phenol-chloroform extraction and precipitation with ethanol as described^1^. Subsequently, the HMW DNA was dissolved in 50 ml TE (pH 8,0) and precipitated by the addition of 5 ml 3 M sodium acetate (pH 5,2) and 100 ml ice-cold ethanol. The suspension was mixed by slow circular movements resulting in the formation of a white precipitate (HMW DNA), which was collected using a wide-bore 5 ml pipette tip and transferred for 30 sec into a tube containing 5 ml 75% ethanol. The washing was repeated twice. The HMW DNA was transferred into a 2 ml tube using a wide-bore tip, collected with a polystyrene spatula, air-dried in a fresh 2 ml tube and dissolved in 500 µl 10 mM Tris-Cl (pH 8.0). For quantification the Qubit dsDNA High Sensitivity assay kit (Thermo Fisher Scientific, MA, USA) was used. The DNA size-profile was recorded using the Femto Pulse system and the Genomic DNA 165 kb kit (Agilent Technologies Inc, CA, USA). In typical experiments the peak of the size-profile of the HMW DNA for library preparation was around 165 kb.

## DNA library preparation and Pacbio HiFi sequencing

For fragmentation of the HMW DNA into 20 kb fragments, a Megaruptor 3 device (speed: 30) was used (Diagenode, NJ, USA). A minimum of two HiFi SMRTbell libraries were prepared for each barley genotype following essentially the manufacturer’s instructions and the SMRTbell Express Template Prep Kit (Pacific Biosciences, CA, USA). The final HiFi libraries were size selected (narrow size range: 18-21 kb) using the SageELF system with a 0,75% Agarose Gel Cassette (Sage Sciences, MA, USA) according to standard manufacturer’s protocols.

HiFi CCS reads were generated operating the PacBio Sequel IIe instrument (Pacific Biosciences, CA, USA) following the manufacturer’s instructions. Per genotype about four 8 M SMRT cells (average yield: 24 Gb HiFi CCS per 8 M SMART cell) were sequenced to obtain an approximate haploid genome coverage of about 20-fold. In typical experiments the concentration of the HiFi library on plate was 80-95 pM. 30 h movie time, 2 h pre-extension and sequencing chemistry v2.0 were used. The resulting raw data was processed using the CCS4 algorithm (https://github.com/PacificBiosciences/ccs).

## Hi-C library preparation and Illumina sequencing

*In situ* Hi-C libraries were prepared from one-week old barley seedlings based on the previously published protocol^2^. Dovetail Omni-C data were generated for Bowman, Aizu6, Golden Melon, 10TJ18 as per manufacturer’s instructions (https://dovetailgenomics.com/products/omni-c-product-page/). Sequencing and Hi-C raw data processing was performed as described before^3,4^.

## Genome sequence assembly and validation

PacBio HiFi reads were assembled using hifiasm (v0.11-r302) ^5^. Pseudomolecule construction was done with the TRITEX pipeline^6^. Chimeric contigs and orientation errors were identified through manual inspection of Hi-C contact matrices. Genome completeness and consensus accuracy were evaluated using Merqury (v1.3) ^7^. Levels of duplication and heterozygosity were assessed with Merqury and FindGSE (v1.94) ^8^. BUSCO (Benchmarking Universal Single-Copy Orthologs) (v.3.0.2) ^9^ with the embryophyta_odb9 data set was run on the final assemblies.

## Single-copy pangenome construction

The single-copy regions in each chromosome-level assembly were identified by filtering 31-mers occurring more than once in the genomic regions by BBDuk (BBMap_37.93, https://jgi.doe.gov/data-and-tools/software-tools/bbtools). Then, the single-copy regions were obtained in BED format and their sequences were retrieved using BEDTools (v2.29.2) ^10^. The single-copy sequences were clustered using MMseqs2 (Many-against-Many sequence searching) ^11^ with the parameters “–-cluster-mode” and setting over 95% sequence identity. A representative from each cluster (the largest in a cluster) was selected to estimate the pangenome size.

## Illumina resequencing

A total of 1,000 plant genetic resources and 315 elite barley varieties (**Supplementary Table 5**) were used for whole genome resequencing. Illumina Nextera libraries were prepared and sequenced on an Illumina NovaSeq 6000 at IPK Gatersleben (**Supplementary Table 5**).

## SNP and SV calling

Reciprocal genome alignment in which each of the pangenome assemblies was aligned to the MorexV3 assembly with the latter acting either as alignment query or reference. Alignment was done with Minimap2 (version 2.20) ^12^. From the resultant two alignment tables, insertion and deletions were called by Assemblytics (version 1.2.1) ^13^ and only deletions were selected in both alignments to convert into presence/absence variants relative to the Morex reference genome. Further, balanced rearrangements (inversions, translocations) were scanned for with SyRI^14^. To call SNPs, raw sequencing reads were trimmed using cutadapt (version 3.3) ^15^ and aligned to the MorexV3 reference genome using Minimap2 (version 2.20) ^12^. The resulting alignments were sorted with Novosort (V3.09.01) (http://www.novocraft.com). BCFtools (version 1.9) ^16^ was used to call SNPs and short insertions and deletions. GWAS for was done with GEMMA^17^.

## Preparation and Illumina sequencing of narrow-size WGS libraries for core50

10 µg DNA in 130 µl were sheared in tubes (Covaris microTUBE AFA Fiber Pre-Slit Snap Cap) to an average size of approximately 250 bp using a Covaris S220 focused-ultrasonicator (peak incidence power: 175 W, duty factor: 10%; cycles per burst: 200; time: 180 sec) according to standard manufacturer’s protocols (Covaris Ltd., Brighton, UK). The sheared DNA was size selected using a BluePippin device and a 1.5% agarose cassette with internal R2 marker (Sage Sciences, MA, USA). A tight size setting at 260 bp was used for the purification of fragments in the narrow range between 200-300 bp (typical yield: 1-3 µg). The size selected DNA was used for the preparation of PCR-free whole genome shotgun libraries using the Roche KAPA Hyper Prep kit according to the manufacturer’s protocols (Roche Diagnostics Deutschland GmbH,

Mannheim, Germany). A total of 10 to 12 libraries were provided with unique barcodes, pooled at equimolar concentrations and quantified by qPCR using the KAPA Library Quantification Kit for Illumina Platforms according to standard protocols (Roche Diagnostics Deutschland GmbH, Mannheim, Germany). The pools were sequenced (2x 151 bp, paired-end) using four S4 XP flowcells and the Illumina NovaSeq 6000 system (Illumina Inc., San Diego, CA, USA) at IPK Gatersleben.

## Contig assembly of core50 sequencing data

Raw reads were demultiplexed based on index sequences and duplicate reads were removed from the sequencing data using Fastuniq^18^. The read1 and read2 sequences were merged based on the overlap using bbmerge.sh from bbmap (v37.28) ^19^. The merged reads were error corrected using BFC (v181) ^20^. The error corrected merged reads were used as an input for Minia3 (v3.2.0) ^21^ to assemble reads into unitigs with the following parameters, –no-bulge-removal –no-tip-removal –no-ec-removal –out-compress 9 –debloom original. The Minia3 source was assembled to enable k-mer size up to 512 as described in the Minia3 manual. Iterative Minia3 runs with increasing k-mer sizes (100, 150, 200, 250 and 300) were used for assembly generation as provided in the GATB Minia pipeline (https://github.com/GATB/gatb-minia-pipeline). In the first iteration, k-mer size of 50 was used to assemble input reads into unitigs. In the next runs, the input reads as well as the assembly of the previous iteration were used as input for the minia3 assembler. BUSCO analysis was conducted on the contig assemblies using BUSCO (v.3.0.2) with embryophyta_odb9 data set^9^. In addition, high-confidence gene models from Morex V3 reference^22^ were aligned to the contig assemblies to assess completeness with the parameter of ≥90% query coverage and ≥97% identity.

## Pangenome accession in diversity space

Pseudo-FASTQ paired-end reads (10-fold coverage) were generated from the 76 pangenome assemblies with fastq_generator (https://github.com/johanzi/fastq_generator) and aligned to MorexV3 reference genome sequence assembly^22^ using Minimap2 (version 2.24-r1122, ref. ^12^). SNPs were called together with short-read data (**Supplementary Table 5**) using BCFtools^23^ version 1.9 with the command “mpileup –q 20 –Q20 –-excl-flags 3332”. To plot the diversity space of cultivated barley, the resultant variant matrix was merged with that of 19,778 domesticated barleys of Milner et al. ^24^ (genotyping-by-sequencing [GBS] data). SNPs with more than 20 % missing or more than 20 % heterozygous calls were discarded. PCA was done with smartpca^25^ version 7.2.1. To represent the diversity of wild barleys, we used published GBS and whole-genome sequencing (WGS) data of 412 accessions of that taxon^26,27^. Variant calling for GBS data was done with BCFtools^23^ (version 1.9) using the command “mpileup –q 20 –Q20”. The resultant variant matrix was filtered as follows: (1) only bi-allelic SNP sites were kept; (2) homozygous genotype calls were retained if their read depth was ≥ 2 and ≤ 50 and set to missing otherwise; (3) heterozygous genotype calls were retained if the read depth of both alleles was ≥ 2 and set to missing otherwise. SNPs with more than 20 % missing, more than 20 % heterozygous calls or a minor allele frequency below 5 % were discarded. PCA was done with smartpca^25^ version 7.2.1. A matrix of pairwise genetic distances based on identity-by-state (IBS) was computed with Plink2 (version 2.00a3.3LM, ref. ^28^) and used to construct an NJ tree with Fneighbor (http://emboss.toulouse.inra.fr/cgi-bin/emboss/fneighbor) in the EMBOSS package^29^. The tree was visualized with Interactive Tree Of Life (iTOL) ^30^.

## Haplotype representation

Pangenome assemblies were mapped to MorexV3 as described above (“Pangenome accession in diversity space”). Read depth was calculated with SAMtools^23^ version 1.16.1. Genotype calls were set to missing if they were supported by fewer than two reads. Identity-by-state (IBS) was calculated with PIink2 (version 2.000a3.3LM, ref. ^28^) in 1 Mb windows (shift: 0.5 Mb) using the using command “--sample-diff counts-only counts-cols=ibs0, ibs1”. Windows which in one of both accessions in the comparison had 2-fold coverage over less than 200 kb were set to missing. The number of differences (d) in a window was calculated as ibs0+ibs1/2, where ibs0 is the number of homozygous differences and ibs1 that of heterozygous ones. This distance was normalized for coverage by the formula d / i x 1 Mb, where i is the size in bp of the region covered in both accessions in the comparison had at least 2-fold coverage. In each window, we determined for each among the PGR and cultivars panel, the closest pangenome accession according to the coverage-normalized IBS distance. Only accessions with fewer than 10 % missing windows due to low coverage were considered, leaving 899 PGRs and 264 cultivars. The distance to the closest pangenome accession was plotted with the R package ggplot2 to determine the threshold for similarity (**Extended Data** Fig. 2d).

## Transcriptome sequencing for gene annotation

Data for transcript evidence-based genome annotation was provided by the International Barley Pan-Transcriptome Consortium, and a detailed description of sample preparation and sequencing is provided elsewhere. In brief, the 20 genotypes sequenced for the first version of the barley pangenome^26^ were used for transcriptome sequencing. Five separate tissues were sampled for each genotype. These were: embryo (including mesocotyl and seminal roots), seedling shoot, seedling root, inflorescence and caryopsis. Three biological replicates were sampled from each tissue type, amounting to 330 samples. Four samples failed quality control and were excluded.

Preparation of the strand-specific dUTP RNA-Seq libraries, and Illumina paired-end 150 bp sequencing were carried out by Novogene (UK) Company Limited. In addition, PacBio Iso-Seq sequencing was carried out using a PacBio Sequel IIe sequencer at IPK Gatersleben. For this, a single sample per genotype was obtained by pooling equal amounts of RNA from a single replicate from all five tissues. Each sample was sequenced on an individual 8M SMRT cell.

## De novo gene annotation

Structural gene annotation was done combining de novo gene calling and homology-based approaches with RNAseq, IsoSeq, and protein datasets (**Extended Data** Fig. 3a). Using evidence derived from expression data, RNAseq data were first mapped using STAR^31^ (version 2.7.8a) and subsequently assembled into transcripts by StringTie^32^ (version 2.1.5, parameters –m 150-t –f 0.3). Triticeae protein sequences from available public datasets (UniProt^33^, https://www.uniprot.org, 05/10/2016) were aligned against the genome sequence using GenomeThreader^34^ (version 1.7.1; arguments –startcodon –finalstopcodon –species rice – gcmincoverage 70 –prseedlength 7 –prhdist 4). Isoseq datasets were aligned to the genome assembly using GMAP^35^ (version 2018-07-04). All assembled transcripts from RNAseq, IsoSeq, and aligned protein sequences were combined using Cuffcompare^36^ (version 2.2.1) and subsequently merged with StringTie (version 2.1.5, parameters –-merge –m150) into a pool of candidate transcripts. TransDecoder (version 5.5.0; http://transdecoder.github.io) was used to identify potential open reading frames and to predict protein sequences within the candidate transcript set.

*Ab initio* annotation was initially done using Augustus^37^ (version 3.3.3). GeneMark^38^ (version 4.35) was additionally employed to further improve structural gene annotation. To avoid potential over-prediction, we generated guiding hints using the above described RNAseq, protein, and IsoSeq datasets as described before^39^. A specific Augustus model for barley was built by generating a set of gene models with full support from RNAseq and IsoSeq. Augustus was trained and optimized following a published protocol^39^. All structural gene annotations were joined using EVidenceModeller^40^ (version 1.1.1), and weights were adjusted according to the input source: *ab initio* (Augustus: 5, GeneMark: 2), homology-based (10). Additionally, two rounds of PASA^41^ (version 2.4.1) were run to identify untranslated regions and isoforms using the above described IsoSeq datasets.

We used BLASTP^42^ (ncbi-blast-2.3.0+, parameters –max_target_seqs 1 –evalue 1e–05) to compare potential protein sequences with a trusted set of reference proteins (Uniprot Magnoliophyta, reviewed/Swissprot, downloaded on 3 Aug 2016; https://www.uniprot.org). This differentiated candidates into complete and valid genes, non-coding transcripts, pseudogenes, and transposable elements. In addition, we used PTREP (Release 19; http://botserv2.uzh.ch/kelldata/trep-db/index.html), a database of hypothetical proteins containing deduced amino acid sequences in which internal frameshifts have been removed in many cases. This step is particularly useful for the identification of divergent transposable elements with no significant similarity at the DNA level. Best hits were selected for each predicted protein from each of the three databases. Only hits with an e-value below 10e–10 were considered. Furthermore, functional annotation of all predicted protein sequences was done using the AHRD pipeline (https://github.com/groupschoof/AHRD).

Proteins were further classified into two confidence classes: high and low. Hits with subject coverage (for protein references) or query coverage (transposon database) above 80% were considered significant and protein sequences were classified as high-confidence using the following criteria: protein sequence was complete and had a subject and query coverage above the threshold in the UniMag database or no BLAST hit in UniMag but in UniPoa and not PTREP; a low-confidence protein sequence was incomplete and had a hit in the UniMag or UniPoa database but not in PTREP. Alternatively, it had no hit in UniMag, UniPoa, or PTREP, but the protein sequence was complete. In a second refinement step, low-confidence proteins with an AHRD-score of 3* were promoted to high-confidence.

## Transposon masking for de novo gene detection

The 20 barley accessions with expression data were softmasked for transposons prior to the *de novo* gene detection using the REdat_9.7_Triticeae section of the PGSB transposon library^43^. Vmatch (http://www.vmatch.de) was used as matching tool with the following parameters: identity>=70%, minimal hit length 75 bp, seedlength 12 bp (vmmatch –d –p –l 75 –identity 70 –seedlength 12 –exdrop 5 –qmaskmatch tolower). The percentage masked was around 80% and almost identical for all 20 accessions.

## Gene projections

Gene contents of the remaining 56 barley genotypes were modelled by the projection of high confidence (HC) genes based on evidence-based gene annotations of the 20 barley genotypes described above. The approach was similar to and built upon a previously described method^26^. To reduce computational load, 760,078 HC-genes of the 20 barley annotations were clustered by cd-hit^44^ requiring 100% protein sequence similarity and a maximal size difference of four amino acids. The resulting 223,182 source genes were subsequently used for all downstream projections as non-redundant transcript set representative for the evidence-based annotations. For each source, its maximal attainable score was determined by global protein self-alignment using the Needleman-Wunsch algorithm as implemented in Biopython^45^ v1.8 and the blosum62 substitution matrix^46^ with a gap open and extension penalty of 0.5 and 10.0, respectively.

Next, we surveyed each barley genome sequence using minimap2 (ref. ^12^) with options ‘-ax splice:hq’ and ‘-uf’ for genomic matches of source transcripts. Each match was scored by its pairwise protein alignment with the source sequence that triggered the match. Only complete matches with start and stop codons and a score ≥0.9 of the source self-score (see above) were retained. The source models were classified into four bins by decreasing confidence qualities: with or without pfam domains, plastid-and transposon-related genes. Projections were performed stepwise for the four qualities, starting from the highest to the lowest. In each quality group, matches were then added into the projected annotation if they did not overlap with any previously inserted model by their coding region. Insertion order progressed from the top to the lowest scoring match. In addition, we tracked the number of insertions for each source by its identifier. For the two top quality categories, we performed two rounds of projections, firstly inserting each source maximally only once followed by rounds allowing one source inserted multiple times into the projected annotation. To consolidate the 20 evidence-based, initial annotations for any genes potentially missed, we employed an identical approach but inserted any non-overlapping matches starting from the prior RNA-seq based annotation. Phylogenetic Hierarchical Orthogroups (HOGs) based on the primary protein sequences from 76 annotated barley genotypes were calculated using Orthofinder^47^ version 2.5.5 (standard parameters). Conserved HOGs contain at least one gene model from all 76 barley genotypes. Variable HOGs contain gene models from at least one barley genotypes and at most 75 barley genotypes. The distribution of all HOG configurations is provided in **Extended Data** Fig. 3b. GENESPACE^48^ was used to determine syntenic relationships between the chromosomes of all 76 genotypes.

## Whole-genome pangenome graphs

Genome graphs were constructed using Minigraph^49^ version 0.20-r559. Other graph construction tools (PGGB^50^, Minigraph-Cactus^51^) turned out to be computationally prohibitive for a genome of this size and complexity, combined with the large number of accessions used in this investigation. Minigraph does not support small variants (< 50 bp), thus graph complexity is lower than with other tools. However, even with Minigraph, graph construction at the whole genome level was computationally prohibitive and thus graphs had to be computed separately for each chromosome, precluding detection of interchromosomal translocations.

Graph construction was initiated using the Morex V3 assembly^52^ as a reference. The remaining assemblies were added into the graph sequentially, in order of descending dissimilarity to Morex. Structural variants were called after each iteration using gfatools bubble (v. 0.5-r250-dirty, https://github.com/lh3/gfatools). Following graph construction, the input sequences of all accessions were mapped back to the graph using Minigraph with the “--call” option enabled, which generates a path through the graph for each accession. The resulting BED format files were merged using Minigraph’s mgutils.js utility script to convert them to P lines and then combined with the primary output of Minigraph in the proprietary RGFA format (https://github.com/lh3/gfatools/blob/master/doc/rGFA.md). Graphs were then converted from RGFA format to GFA format (https://github.com/GFA-spec/GFA-spec/blob/master/GFA1.md) using the “convert” command from the vg toolkit^53^ version v1.46.0 “Altamura”. This step ensures that graphs are compatible with the wider universe of graph processing tools, most of which require GFA format as input. Chromosome-level graphs were then joined into a whole-genome graph using vg combine. The combined graph was indexed using vg index and vg gbwt, two components of the the vg toolkit^53^.

General statistics for the whole-genome graph were computed with vg stats. Graph growth was computed using the heaps command from the ODGI toolkit^54^ version 0.8.2-0-g8715c55, followed by plotting with its companion script heaps_fit.R. The latter also computes values for gamma, the slope coefficient of Heap’s law which allows the classification of pangenome graphs into open or closed pangenomes, i.e. a prediction of whether the addition of further accessions would increase the size of the pangenome^55^.

Structural variant (SV) statistics were computed based on the final BED file produced after the addition of the last line to the graph. A custom shell script was used to classify variants according to the Minigraph custom output format. This allows the extraction of simple, i.e. non-nested, insertions and deletions (relative to the MorexV3 graph backbone), as well simple inversions. The remaining SVs fall into the “complex” category where there can be multiple levels of nesting of different variant types and this precluded further, more fine-grained classification.

To elucidate the effect of a graph-based reference on short read mapping, we obtained whole genome shotgun Illumina reads from five barley samples (**Extended Data** Fig. 4b) in the European Nucleotide Archive (ENA) and mapped these onto the whole genome graph using vg giraffe^56^. For comparison with the standard approach of mapping reads to a linear single genome reference, we mapped the same reads to the Morex V3 reference genome sequence assembly^52^ with bwa mem^57^ version 0.7.17-r1188. Mapping statistics were computed with vg^53^ stats and samtools^23^ stats (version 1.9), respectively.

## Analysis of the *Mla* locus

The coordinates and sequences of the 32 genes present at the *Mla* locus were extracted from the MorexV3 genome sequence assembly^52^. To find the corresponding position and copy number in each of the 76 genomes, we used BLAST^42^ (-perc_identity: 90, –word_size:11, all other parameters set as default). The expected BLAST result for a perfectly conserved allele is a long fragment (exon_1) of 2,015 bp follow by a gap of ∼1,000 bp due to the intron and another fragment (exon_2) of 820 bp. To detect the number of copies, first multiple BLAST results for a single gene were merged if two different BLAST segments were within 1.1kb. Then only if the total length of the input was found, this was counted as a copy. To analyse the structural variation across all 76 accessions, the non-filtered BLAST results were plotted in a region of –20,000 and +500,000 base pairs around the start of the BPM gene HORVU.MOREX.r3.1HG0004540 that was used as an anchor (present in all 76 lines, Supplementary Figure 1. To detection the different *Mla* alleles, three different threshold of – Perc_identity for the BLAST were used: 100, 99 and 98.

## Scan for structurally complex loci

We utilised a pipeline developed by Rabanus-Wallace et al. ^58^ that performs sequence-agnostic identification of long-duplication-prone-regions (l-DPRs) in a reference genome, followed by identification of gene families with a statistical tendency to occur within l-DPRs. The pipeline assumes that a candidate l-DPR will contain an elevated concentration of locally repeated sequences in the kb-scale length range. We first aligned the MorexV3 genome sequence assembly^52^ against itself using lastz^59^ (v1.04.03; arguments: ‘--notransition –-step=500 –gapped’). For practicality purposes, this was done in 2 Mb blocks with a 200 kb overlap, and any overlapping l-DPRs identified in multiple windows were merged. For each window, we ignored the trivial end-to-end alignment, and of the remaining alignments, retained only those longer than 5 kb and falling fully within 200 kb of one and another. An alignment ‘density’ was calculated over the chromosome by calculating, at ‘interrogation points’ spaced equally at 1 kb intervals along the length of the chromosome, an alignment density score that is simply the sum of all the lengths of any of the filtered alignments spanning that interrogation point. A Gaussian kernel density (bandwidth 10 kbp) was calculated over these the interrogation points, weighted by their scores. To allow comparability between windows, the interrogation point densities were normalised by the sum of scores in the window. Runs of interrogation points at which the density surpassed a minimum density threshold were flagged as l-DPRs. A few minor adjustments to these regions (merging of overlapping regions, and trimming the end coordinates to ensure the stretches always begin and end in repeated sequence) yielded the final tabulated list of l-DPR coordinates (**Supplementary Table 7**). The method was implemented in R making use of the package data.table. Genes in each l-DPRs were clustered with UCLUST^60^ (v11, default parameters) using a protein clustering distance cutoff of 0.5 and for each cluster the most frequent functional description as per the MorexV3 gene annotation^52^ was assigned as the functional description of the cluster.

## Molecular dating of divergence times of duplicated genes in complex loci

For molecular dating of gene duplications, we used segments of up to 4 kb, starting 1 kb upstream of duplicated genes in complex loci. With this, we presumed to only use intergenic sequences which are free from selection pressure and thus evolve at a neutral rate of 1.3×10^-8^ substitutions per site per year^61^. The upstream sequences of all duplicated genes of respective complex locus were then aligned pairwise with the program Water from the EMBOSS package^29^ (obtained from Ubuntu repositories, ubuntu.com). This was done for all gene copies of all barley accession for which multiple gene copies were found. Molecular dating of the pairwise alignments was done as previously described^62^ using the substitution rate of 1.3×10^-8^ substitutions per site per year^61^.

## *Amy1_1* analysis in pangenome assemblies

The *amy1_1* gene copy HORVU.MOREX.PROJ.6HG00545380 was used was used to BLAST against all 76 genome assemblies. Full-length sequences with identity over 95% were extracted and used for further analyses. Unique sequences were identified by clustering at 100% identity using CD-Hit^44^ and were aligned using MAFFT^63^ v7.490. Sequence variants among *amy1_1* gene copies at genomic DNA, CDS and respective protein level were collected and *amy1_1* haplotypes (i.e. the combinations of copies) in each genotype assembly were summarized using R^64^ v4.2.2. A Barke-specific SNP locus (GGCGCCAGGCATGATCGGGTGGTGGCCAGCCAAGGCGGTGACCTTCGTGGACAACCACGACACCGGCTCCACGCAGCACATGTGGCCCTTCCCTTCTGACA[A/G]GGTCATGCAGGGATATGCGTACATACTCACGCACCCAGGGACGCCATGCATCGTGAGTTCGTCGTACCAATACATCACATCTCAATTTTCTTTTCTTGTTTCGTTCATAA) for *amy1_1* haplotype cluster ProtHap3 (**Supplementary Table 20**) was identified and used for KASP marker development (LGC Biosearch Technologies, Hoddesdon, United Kingdom).

## Comparative analysis of the *amy1_1* locus structure

Based on the genome annotation of cv. Morex, 15 gene sequences on either side of *amy1_1* gene copy HORVU.MOREX.PROJ.6HG00545440 were extracted. The 31 genes were compared against the 76 genome assemblies using NCBI-BLAST^42^ (BLASTN, word_size of 11 and percent identity of 90, other parameters as default). Alignment plots were generated from the BLAST result coordinates by scaling based on the mid-point between HORVU.MOREX.r3.6HG0617300/HORVU.MOREX.PROJ.6HG00545250 and HORVU.MOREX.r3.6HG0617710/HORVU.MOREX.PROJ.6HG00545670. All BLAST results in the region (+/-1Mb) around this mid-point were plotted using R^64^.

## *Amy1_1* PacBio amplicon sequencing

Genomic DNA from one-week old Morex seedling leaves was extracted with DNeasy® Plant Mini Kit (QIAGEN GmbH, Hilden, Germany). Based on the MorexV3 genome sequence assembly^52^, *amy1_1* full-length copy-specific primers were designed using Primer3 (ref. ^65^) (https://primer3.ut.ee/): 6F – GTAGCAGTGCAGCGTGAAGTC, 80F – AGACATCGTTAACCACACATGC, 82F – GTTTCTCGTCCCTTTGCCTTAA, 82F – GTTTCTCGTCCCTTTGCCTTAA, 33R – GATCTGGATCGAAGGAGGGC, 79R – TCATACATGGGACCAGATCGAG, 80R – ACGTCAAGTTAGTAGGTAGCCC. All forward primers were tagged with bridge sequence (preceding T to primer name) [AmC6]gcagtcgaacatgtagctgactcaggtcac, while reverse primers were tagged with [AmC6]tggatcacttgtgcaagcatcacatcgtag to allow annealing to barcoding primers. These bridge sequence-tagged gene-specific primers were used in pairs with each other targeting 1-2 copies of 3-6 kb *amy1_1* genes, including upstream and downstream 500-1000 bp regions: T6F + T33R, T6F + T79R, T80F + T80R and T82F + T80R. A two-step PCR protocol was conducted. The first step PCR reaction was prepared in 25 μl volume using 2 μl DMSO, 0.3 μl Q5 polymerase (New England Biolab, Massachusetts, United States), 1 μl amy1_1-specific primer pair (10 μM each), 2 μl gDNA, 0.5 μl dNTPs (10mM), 5 μl Q5 buffer and H_2_O. The PCR program was as following: initial denaturation at 98°C/1min followed by 25-28 cycles of 98°C/30 sec, 58°C/30 sec, and 72°C/3 min for extension, with a final extension step of 72°C/2 min. The second PCR step (barcoding PCR) was prepared in the same way using 1 μl of the first PCR product as DNA template, barcoding primers (Pacific Biosciences of California, Inc., California, United States) and the PCR program reduced to 20 cycles. After quality check on 1% agarose gel, all barcoded PCR products were mixed and purified with AMPure® PB (Pacific Biosciences of California, Inc., California, United States). The SMRT bell library preparation and sequencing were carried out at BGI Tech Solutions (BGI Tech Solutions Co., Ltd., Hongkong, China). Sequencing data was analysed using SMRT Link v.10.2. To minimize PCR chimeric noise, CCS were first constructed for each molecule. Secondly, Long Amplicon Analysis (LAA) was carried out based on subreads from 50 bp window spanning peak positions of all CCS length. Final consensus sequences for each *amy1_1* was determined with the aid of size estimation from agarose gel imaging.

## *Amy1_1* SNP haplotype analysis and k-mer based copy number estimation

SNP haplotypes were analyzed in 1,315 plant genetic resources and elite varieties in the extended amy1_1-cluster region (MorexV3 chr6H: 516,385,490 – 517,116,415 bp). SNPs with >20% missing data among the analyzed lines and minor allele frequency (MAF) < 0.01 were removed from downstream analyses. The data was converted to 0,1 and 2 format using VCFtools^66^ and samples were clustered using pheatmap package (https://cran.r-project.org/web/packages/pheatmap/pheatmap.pdf) from R statistical environment^58^. The sequential clustering approach was used to achieve the desired separation. At each step, two extreme clusters were selected and then samples from each cluster were clustered separately. The process was repeated until the desired separation was achieved based on visual inspection.

K-mers (k=21) were generated from Morex *amy1_1* gene family member’s conserved region using jellyfish^67^ v2.2.10. After removing k-mers with counts from regions other than *amy1_1* in the Morex V3 genome assembly, k-mers were counted in the Illumina raw reads (**Supplementary Table 5)** using Seal (BBtools, https://jgi.doe.gov/data-and-tools/software-tools/bbtools/). All k-mer counts were normalized to counts per MorexV3 genome and *amy1_1* copy number was estimated as the median count of all k-mers from each accession in R.

Estimation ability was validated by comparing copy number from pangenome assemblies and short-read sequencing data (**Extended Data** Fig. 8c). For 1,000 plant genetic resources, countries (>=10 accessions) were color shaded based on their proportions of accessions with *amy1_1* copy number > 5 on a world map using the R package maptools (https://cran.r-project.org/web/packages/maptools/index.html).

## AMY1_1 protein structure and protein folding simulation

The published protein structure of α-amylase AMY1_1 from accession Menuet, in complex with the pseudo-tetrasaccharide acarbose (PDB:1bg9; ref. ^68^), was used to simulate the structural context of the amino acid variants identified in barley accessions Morex, Barke and RGT Planet. The amino acid sequence of the crystalized AMY1_1 protein from Menuet and the Morex reference copy *amy1_1* HORVU.MOREX.PROJ.6HG00545380 used in this study are identical. The protein was visualized using PyMol 2.5.5 (Schrödinger Inc. New York, NY, USA). The Dynamut2 webserver^69^ was used to predict changes in protein stability and dynamics by introducing amino acid variants identified in the Morex, Barke and RGT Planet genome assemblies.

## Development of diverse *amy1_1* haplotype barley near-isogenic lines

Near-isogenic lines (NILs) with different *amy1_1* haplotypes were derived from crosses between RGT Planet as recipient and Barke or Morex *amy1_1*-cluster donor parents (ProtHap3, ProtHap4 and ProtHap0, respectively; **Supplementary Table 20**), followed by two subsequent backcrosses to RGT Planet and one selfing step (BC_2_S1) to retrieve homozygous plants at the *amy1_1* locus. A total of four amy_1_1-Barke NILs (ProtHap3) and one amy1_1-Morex NIL (ProtHap0) were developed and tested against RGT Planet (ProtHap4) replicates. Plants were grown in a greenhouse at 18°C under 16/8-hour light/dark cycles. Foreground and background molecular markers were used in each generation to assist plant selection. Respective BC_2_S_1_ plants were genotyped with the Barley Illumina 15K array (SGS Institut Fresenius GmbH, TraitGenetics Section, Germany) and grown to maturity. Grains were harvested and further propagated in field plots in consecutive years in various locations (Nørre Aaby, Denmark; Lincoln, New Zealand; Maule, France). Grains from field plots were harvested and threshed using a Wintersteiger Elite plot combiner (Wintersteiger AG, Germany), and sorted by size (threshold, 2.5 mm) using a Pfeuffer SLN3 sample cleaner (Pfeuffer GmbH, Germany).

## Micro-malting and α-amylase activity analysis

Non-dormant barley samples of RGT Planet and respective NILs with different *amy1_1* haplotypes (50g each, graded >2.5 mm) were micro-malted in perforated stainless-steel boxes. The barley samples were steeped at 15 °C by submersion of the boxes in water. Steeping took place for six hours on day one, three hours on day two and one hour on day three, followed by air rests, to reach 35%, 40% and 45% water content, respectively. The actual water uptake of individual samples was determined as the weight difference between initial water content, measured with Foss 1241 NIT instrument (Foss A/S, Hillerød, Denmark), and the sample weight after surface water removal. During air rest, metal beakers were placed into a germination box at 15°C. Following the last steep, the barley samples were germinated for 3 days at 15°C. Finally, barley samples were kiln dried in an MMK Curio kiln (Curio Group Ltd, Buckingham, England) using a two-step ramping profile. First ramping step started at a set point of 27°C and a linear ramping at 2°C/h to the breakpoint at 55°C using 100% fresh air. Second linear ramping was at 4°C/h reaching a maximum at 85°C. This temperature was kept constant for 90 minutes using 50% air recirculation. The kilned samples were then deculmed using a manual root removal system (Wissenschaftliche Station für Brauerei, Munich, Germany). α-amylase activity was measured using the Ceralpha method (Ceralpha Method MR-CAAR4, Megazyme) modified for Gallery Plus Beermaster (Thermo Fisher Scientific, USA).

## Rachilla hair ploidy measurements

Ploidy assessment was performed on rachillae harvested from barley spikes at developmental stage^70^ ∼W9.0. Once isolated, rachillae were fixed with 50% Ethanol/10% acetic acid for 16h after which they were stained with 1 µM 4’,6-Diamidino-2-phenylindol (DAPI) in 50 mM phosphate buffer (pH 7.2) supplemented with 0.05% Triton X100. Probes were analyzed with a Zeiss LSM780 confocal laser scanning microscope using a 20x NA 0.8 objective, zoom 4x, and image size 512 x 512 pixel. DAPI was visualized with a 405 nm laser line in combination with a 405–475 nm bandpass filter. Pinhole was set to ensure the whole nucleus was measured in one scan. Size and fluorescence intensity of the nuclei were measured with ZEN black (ZEISS) software. For data normalization small round nuclei of the epidermal proper were used for 2C calibration.

## Scanning electron microscopy

Sample preparation and recording by scanning electron microscopy was essentially performed as described previously^71^. In brief, samples were fixed overnight at 4°C in 50 mM phosphate buffer (pH 7.2) containing 2% v/v glutaraldehyde and 2% v/v formaldehyde. After washing with distilled water and dehydration in an ascending ethanol series, samples were critical point-dried in a Bal-Tec critical point dryer (Leica microsystems, https://www.leica-microsystems.com). Dried specimens were attached to carbon-coated aluminium sample blocks and gold-coated in an Edwards S150B sputter coater (Edwards High Vacuum Inc., http://www.edwardsvacuum.com). Probes were examined in a Zeiss Gemini30 scanning electron microscope (Carl Zeiss Microscopy GmbH, https://www.zeiss.de) at 5 kV acceleration voltage. Images were digitally recorded.

**Linkage mapping of *SHORT RACHILLA HAIR 1 (HvSRH1)***

Initial linkage mapping was performed using genotyping-by-sequencing (GBS) data of a large ‘Morex’ x ‘Barke’ F_8_ RIL population^72^ (ENA project PRJEB14130). The GBS data of 163 RILs, phenotyped for rachilla hair in the F_11_ generation, and the two parental genotypes were extracted from the variant matrix using VCFtools^66^ and filtered as described previously^24^ for a minimum depth of sequencing to accept heterozygous and homozygous calls of 4 and 6, respectively, a minimum mapping quality score of the SNPs of 30, a minimal fraction of homozygous calls of 30 %, and a maximum fraction of missing data of 25%. The linkage map was built with the R package ASMap^73^ using the MSTMap algorithm^74^ and the Kosambi mapping function, forcing the linkage group to split according to the physical chromosomes. The linkage mapping was done with R/qtl^75^ using the binary model of the scanone function with the EM method^76^. The significance threshold was calculated running 1000 permutations and the interval was determined by a LOD drop of 1. To confirm consistency between the F_8_ RIL genotypes and F_11_ RIL phenotypes, three PCR Allele Competitive Extension (PACE) markers were designed though 3CR Bioscience (Essex, UK) free assay design service, using polymorphisms between the genome assemblies of the two parents (**Supplementary Table 23**), and PACE genotyping was performed as described earlier^77^. To reduce the *Srh1* interval, 22 recombinant F_8_ RILs were sequenced by Illumina whole-genome sequencing (WGS), the sequencing reads were mapped on MorexV3 reference genome^52^, and the SNP called. The 100 bp region around the flanking SNPs of the *Srh1* interval as well as the sequence of the candidate gene HORVU.MOREX.r3.5HG0492730 were compared to the pangenome assemblies using BLASTN^78^ to identify the corresponding coordinates and extract the respective intervals for comparison. Gene sequences were aligned with Muscle5 (ref. ^79^). Structural variation between intervals was assessed with LASTZ^59^ version 1.04.03. The motif search was carried out with the EMBOSS^80^ 6.5.7 tool fuzznuc.

## Cas9-mediated mutagenesis

Guide RNA (gRNA) target motifs in the ‘Golden Promise’ *HvSrh1* candidate gene HORVU.GOLDEN_PROMISE.PROJ.5HG00440000.1 were selected by using the online tool WU-CRISPR^81^ to induce translational frameshift mutations by insertion/deletion of nucleotides leading to loss-of-function of the gene. One pair of target motifs (gRNA1a: CCTCGCTGCCCGCCGACGC, gRNA1b: GACAAGACGAAGGCCGCGG) was selected within the *HvSrh1* candidate gene based on their position within the first half of the coding sequence and the two-dimensional minimum free energy structures of the cognate single-gRNAs (NNNNNNNNNNNNNNNNNNNNGUUUUAGAGCUAGAAAUAGCAAGUUAAAAUAAGGCUAGUCCGUUAUCAACUUGAAAAAGUGGCACCGAGUCGGUGCUUUU) as modelled by the RNAfold WebServer^82^ and validated as suggested by Koeppel et al. ^83^. gRNA-containing transformation vectors were cloned using the modular CasCADE vector system (https://doi.org/10.15488/13200). gRNA-specific sequences were ordered as DNA oligonucleotides (**Supplementary Table 24**) with specific overhangs for BsaI-based cloning into the gRNA-module vectors carrying the gRNA scaffold, driven by the *Triticum aestivum* U6 promoter. Golden Gate assembly of gRNAs and the *cas9* module, driven by the *Zea mays Polyubiquitin 1* (*ZmUbi1*) promotor, were performed according to the CasCADE protocol to generate the intermediate vector pHP21. To generate the binary vector pHP22, the gRNA and *cas9* expression units were cloned using SfiI into the generic vector^84^ p6i-2×35S-TE9 that harbours an *hpt* gene under control of a doubled-enhanced *CaMV35S* promoter in its transfer-DNA for plant selection. *Agrobacterium*-mediated DNA transfer to immature embryos of the spring barley Golden Promise was performed as previously described^85^. In brief, immature embryos were excised from caryopses 12-14 days after pollination and co-cultivated with *Agrobacterium* strain AGL1 carrying pHP22 for 48 hours. Then, the explants were cultivated for further callus formation under selective conditions using Timentin and hygromycin, which was followed by plant regeneration. The presence of T-DNA in regenerated plantlets was confirmed by *hpt*-and *cas9*-specific PCRs (primer sequences in **Supplementary Table 24**). Primary mutant plants (M_1_ generation) were identified by PCR amplification of the target region (primer sequences in **Supplementary Table 24**) followed by Sanger sequencing at LGC Genomics GmbH (Berlin, Germany). Double or multiple peaks in the sequence chromatogram starting around the Cas9 cleavage site upstream of the target’s protospacer-adjacent motif (PAM) were considered as an indication for chimeric and/or heterozygous mutants. Mutant plants were grown in a glasshouse until the formation of mature grains. M_2_ plants were grown in a climate chamber under speed breeding conditions (22 h light at 22 °C and 2 h dark at 19°C, adapted from Watson et al. ^86^ and genotyped by Sanger sequencing of PCR amplicons as given above. M_2_ grains were subjected to phenotyping.

## FIND-IT library construction

We constructed a FIND-IT library in cv. ‘Etincel’ (6-row winter malting barley; SECOBRA Recherches) as described in Knudsen et al. ^87^. In short, we induced mutations by incubating 2.5 kg of ‘Etincel’ grain in water overnight at 8°C following an incubation in 0.3 mM NaN_3_ at pH 3.0 for 2 hours at 20°C with continuous application of oxygen. After thoroughly washing with water, the grains were air-dried in a fume hood for 48 hours. Mutagenized grains were sown in fields in Nørre Aaby, Denmark, and harvested in bulk using a Wintersteiger Elite plot combiner (Wintersteiger AG, Germany). In the following generation, 2.5 kg of grain was sown in fields in Lincoln, New Zealand, and 188 pools of approximately 300 plants each were hand-harvested and threshed. A representative sample, 25% of each pool, was milled (Retsch GM200, Haan, Germany), and DNA was extracted from 25 g of the flour by LGC Genomics GmbH (Berlin, Germany).

## FIND-IT screening

The FIND-IT ‘Etincel’ library was screened as described in Knudsen et al. ^87^ using a single assay for the isolation of *srh1*^P63S^ variant [ID# CB-FINDit-Hv-014]. Forward primer 5’ AATCCTGCAGTCCTTGG 3’, reverse primer 5’ GAGGAGAAGAAGGAGCC 3’, mutant probe 5’6-FAM/CGTGGACGT/ZEN/CGACG/3’IABkFQ/ Wild type probe /5’SUN/ACGTGGGCG/ZEN/TCGA/3’IABkFQ/ Integrated DNA Technologies, Inc.

## 4K SNP chip genotyping

Genotyping, including DNA extraction from freeze-dried leaf material, was conducted by TraitGenetics (SGS – TraitGenetics GmbH, Germany). *srh1*^P63S^ mutant, the corresponding wild type ‘Etincel’ and *srh1* pangenome accessions Morex, RGT Planet, HOR 13942, HOR 9043 and HOR 21599 were genotyped for background confirmation. Pairwise genetic distance of individuals was calculated as the average of their per-locus distances^88^ using R package stringdist^89^ (v 0.9.8). Principal Coordinate Analysis (PCoA) was done with R^64^ (v 4.0.2) base function cmdscale based on this genetic distance matrix. The first two PCs were illustrated by ggplot2 (https://ggplot2.tidyverse.org).

## Sanger sequencing

gDNA of the *srh1*^P63S^ variant and ‘Etincel’ was extracted from one-week old seedling leaves (DNeasy, Plant Mini Kit, Qiagen, Hilden, Germany). Genomic DNA fragments for sequencing were amplified by PCR using gene specific primers (forward primer 5’TTGCACGATTCAAATGTGGT 3’, reverse primer 5’ TCACCGGGATCTCTCTGAAT 3’) and Taq DNA Polymerase (NEB) for 35 cycles (initial denaturation at 94°C/3 min followed by 35 cycles of 94°C/45 s, 55°C/60 s, and 72°C/60 s for extension, with a final extension step of 72°C/10 min). PCR products were purified using the NucleoSpin Gel and PCR Clean-Up Kit (Macherey-Nagel GmbH & Co. AG, Düren, Germany) according to the manufacturer’s instructions. Sanger sequencing was done at Eurofins Genomics Germany GmbH using a gene-specific sequencing primer (5’ AGAACGGAGAGGAGAGAAAGAAG 3’).

## RNA preparation, sequencing, and data analysis

Rachilla tissues from two contrast groups Morex (short), Barke (long) and Bowman (long) and BW-NIL-*srh1* (short) were used for RNA sequencing. The rachilla tissues were collected from the central spikelets of the respective genotypes at rachilla hair initiation (RI; Waddington 8.0), and elongation (RE; Waddington 9.5) stages. Total RNA was extracted using TRIzol reagent (Invitrogen) followed by 2-propanol precipitation. Genomic DNA residues were removed with DNase I (NEB, M0303L). High-throughput paired-end sequencing was conducted at Novogene Co., Ltd (Cambridge, UK) with Illumina NovaSeq 6000 PE150 Platform. RNAseq reads were trimmed for adaptor sequences with Trimmomatic^90^ (version 0.39) andt the MorexV3 genome annotation was used as reference to estimate read abundance with Kallisto^91^. The raw read counts were normalized to Transcripts per kilo base per million (TPM) expression levels.

## *mRNA insitu* hybridization

*in situ* hybridization was conducted in longitudinal and cross sections derived from whole spikelet tissues of Bowman and Morex at rachilla hair elongation developmental stage (W9.5) with Hv*SRH1* sense & antisense probes (124 bp). The *in situ* hybridization was performed as described before^92^ with few modifications.

## Code availability

Scripts for pangenome graphs analyses are available at https://github.com/mb47/minigraph-barley. The scripts for calculation of core/shell and cloud genes are deposited to the repository https://github.com/PGSB-HMGU/. The pipeline for identifying structurally complex loci is available at https://github.com/mtrw/DGS.

## Data availability

All the sequence data collected in this study have been deposited at the European Nucleotide Archive (ENA) under BioProjects PRJEB40587, PRJEB57567 and PRJEB58554 (raw data for pangenome assemblies), PRJEB64639 (pan-transcriptome Illumina data), PRJEB64637 (transcriptome Isoseq data), PRJEB53924 (Illumina resequencing data), PRJEB45466-511 (raw data for gene space assemblies), PRJEB65284 (*srh1* transcriptome data). Accession codes for individual genotypes are listed in supplementary tables: **Supplementary Table 1** (pangenome assemblies and associated raw data), **Supplementary Table 2** (transcriptome data), **Supplementary Table 5** (Illumina resequencing), **Supplementary Table 6** (gene space assemblies).

## Notes

### Competing Interest Statement

K.B., C.D., M.E.J., S.M.K., Q.L., E.M., P.R.P., B.Skadhauge., H.C.T., M.T.S.N., C.V., M.W.R. are current or previous Carlsberg A/S employees. P.A.P. and D.V. are SECOBRA Recherches employees. All other authors declare no competing interests.

