## Supplementary Figures for "Adaptive diversification through structural variation in barley"

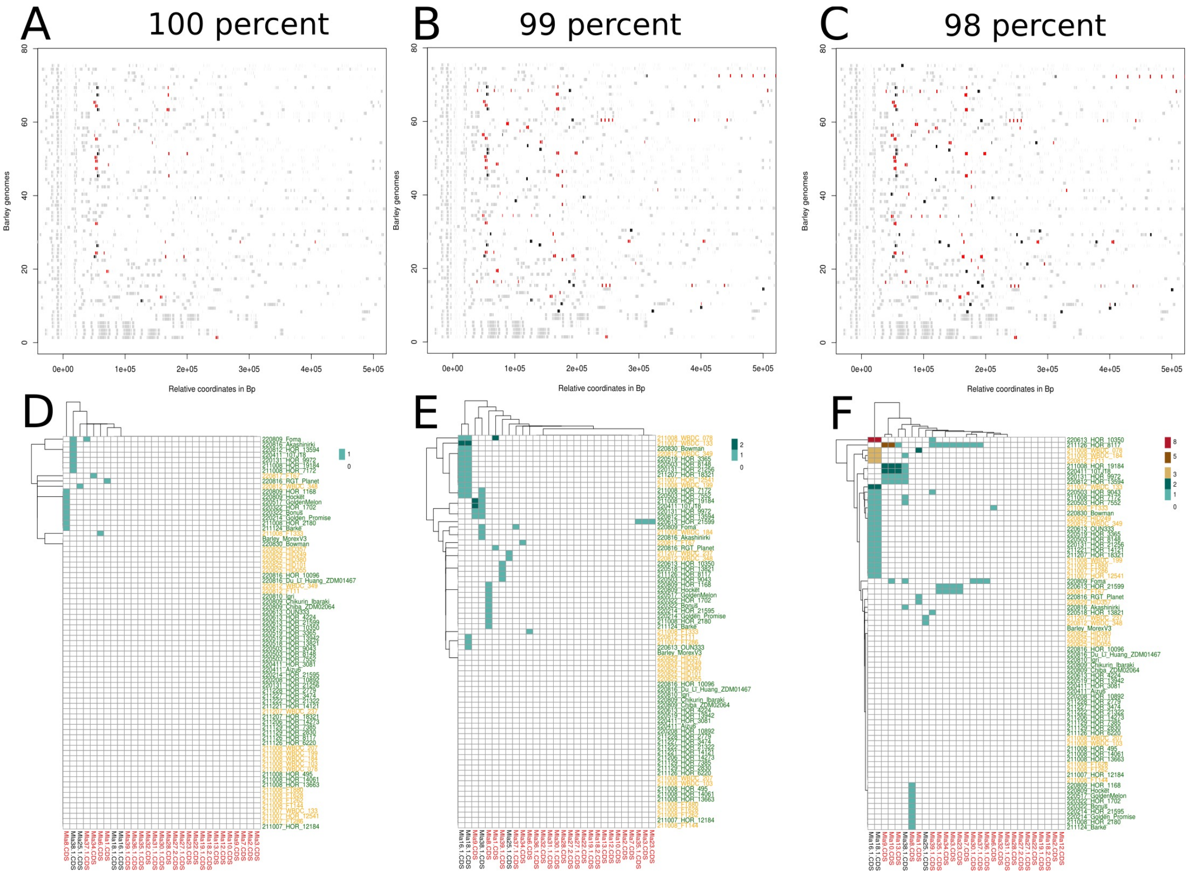


**Supplementary Figure 1: Structure and copy number variation at *Mla* at different thresholds for alignment similarity.** Structural plot around the *Mla* locus for the 76 genomes. The gray rectangles represent all the homologous genes (based on BLAST alignments) in the regions in comparison with the Morex annotation. Each rectangle represents a blast result and the size on the rectangle is proportional to the length of the matching fragment. The red and black rectangles represent the position of the known *Mla* alleles from subfamily 1 and 2, respectively, as defined by Seeholzer et al.[1] Each gene coordinate is scaled based on the position of the *Bpm* gene in the respective genome. Three different thresholds, 100, 99, and 98%, for blasting the *Mla* alleles have been used for **(a)**, **(b)**, and **(c)**, respectively. **(d-f)**: Copy number variation of *Mla* allele in the 76 barley genome. *Mla* alleles names in red and black represent the two subfamilies. The names of the accessions are coloured according to domestication states (green – domesticated; orange – wild). The colouring of the square indicates the number of copies according to the legend.

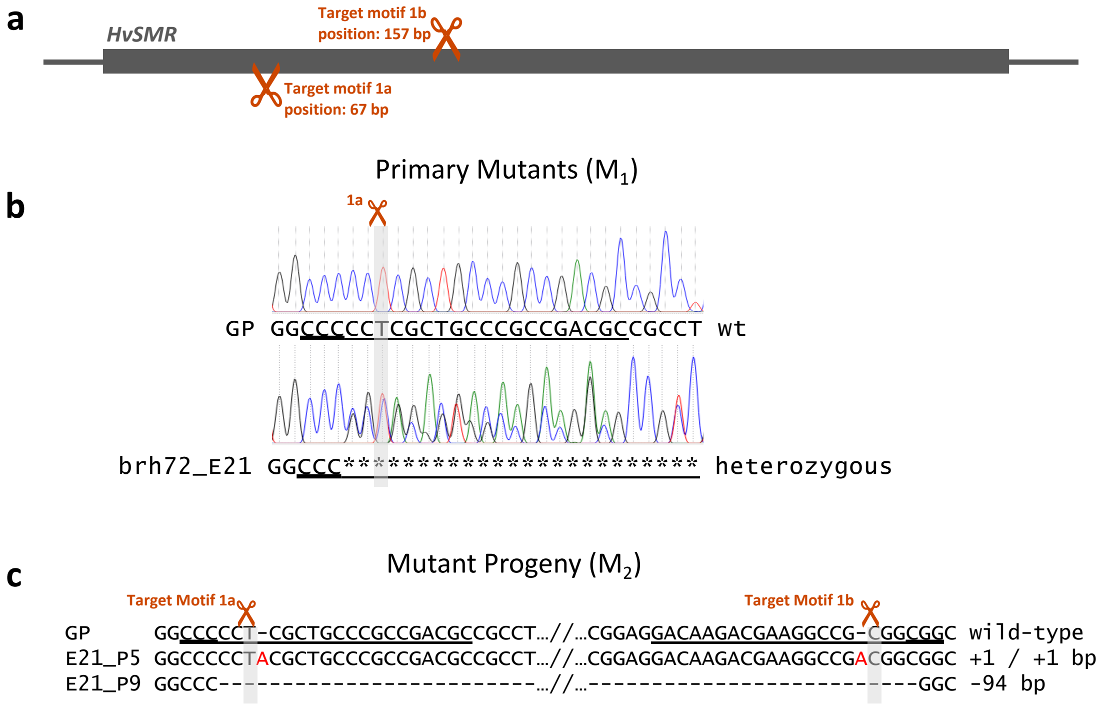


**Supplementary Figure 2: Targeted mutagenesis at *HvSRH1.*** **(a)** Structure of the *srh1* candidate gene with the only exon shown as a grey box and localization of target motifs 1a and 1b in a distance of 87 bp between the expected cleavage sites of Cas9. Target motif 1a is located at the non-coding strand and 1b at the coding strand. **(b)** Chromatogram of Sanger sequencing of PCR amplicons of target motif 1a in Golden Promise (GP) wild-type (wt) and the primary mutant brh72_E21. Double peaks in the chromatogram indicate heterozygous insertions and/ or deletions. **(c)** Homozygous mutations in two individuals of the $M_2$ progeny of plant E21. Dashes indicate deleted nucleotides, red letters indicate insertions, asterisks indicate unclear signal due to double peaks, protospacer adjacent NGG motif doubled underlined, target motif underlined, scissors indicate cleavage site of Cas9.
